# Very Early Cortical Auditory Responses to Speech in Humans

**DOI:** 10.64898/2026.06.04.730183

**Authors:** Karl D. Lerud, Charlie Fisher, Vrishab Commuri, Samira Anderson, Jonathan Z. Simon

**Affiliations:** Institute for Systems Research, University of Maryland, College Park MD 20742; Department of Electrical & Computer Engineering, University of Maryland, College Park MD 20742; Department of Hearing and Speech Sciences, University of Maryland, College Park MD 20742; Brain and Behavior Institute, University of Maryland, College Park MD 20742; Department of Biology, University of Maryland, College Park MD 20742

**Keywords:** auditory, cortex, speech, EEG, MEG, ABR, TRF, MLR, Wave V, SN10

## Abstract

A foundational measure of the auditory system’s ability to process sound correctly is the auditory brainstem response (ABR) and middle latency response (MLR) complex, but their dependence on stimulus complexity and behavioral relevance is poorly understood. Here we recorded noninvasive, simultaneous electroencephalography (EEG) and magnetoencephalography (MEG) in adults listening to both naturalistic speech and clicks, and computed the resulting temporal response functions (TRFs). The TRF’s ABR-MLR complex for speech stimuli contains a very early cortical response peak (11 ms latency), a processing stage which has not been previously well-characterized, and which is absent for responses to clicks. Other significant differences in latency and source between the speech and click responses demonstrate further differences in auditory processing in both thalamus and cortex. In contrast, the speech and click TRF’s ABR complex share a common, prominent wave V response, indicating their similar processing in midbrain. In summation, the use of naturalistic speech as a stimulus reveals how even very early auditory responses reflect its ethological relevance in humans.

## Introduction

The ability of the mammalian auditory system to quickly and efficiently distinguish between relevant and irrelevant sounds in the acoustic environment is essential to survival, and the neural underpinnings of these abilities are a deep and ongoing area of human auditory neuroscience. At the level of auditory cortex, this includes investigations of brain responses corresponding to selective attention in an acoustic environment with distractors,^1–7^ and includes investigations of the neural origins (e.g., latencies and neural sources) of responses to sounds of diverse potential biological relevance, including speech sounds.^8–11^ Both electroencephalography (EEG) and magnetoencephalography (MEG) recordings are used to investigate these types of auditory responses, with EEG being far more common. For subcortical and early cortical responses, the auditory brainstem response (ABR) and middle latency response (MLR) are well-established early neural aspects of auditory responses, as revealed by evoked responses to clicks (and other punctate sounds). However, how they are related to responses to other stimuli, such as the periodic frequency following response (FFR) or auditory steady-state response (ASSR) is still not well understood, and their relationship to more complex sounds such as natural speech is even less well understood. Additionally, while the neural sources of the ABR are well established^12,13^ and the neural sources of the late MLR peaks are dominantly cortical,^14,15^ the neural origins of the early MLR peaks also still remain elusive.

Prominent peaks in the ABR-MLR complex and their associated latencies include wave V (5 – 8 ms), Na (10 – 21 ms), Pa (25 – 30 ms), Nb (30 – 45 ms), and P50 (45 – 60 ms, also referred to as the Pb in the context of the MLR and the P1 in auditory evoked responses). The literature offers some variability as to the precise latency and sources of these six components (wave V, SN10, Na, Pa, Nb, and P50), as well as to which stimuli maximally elicit them, where some of the variability in latency can be ascribed to differences in the eliciting stimuli.^16–20^ Wave V is widely understood to originate in the inferior colliculus.^12,13^ The Na complex, a post-brainstem inflection point of particular interest in this study, is often subdivided into two parts (both characterized by EEG vertex-negative polarity), with corresponding latencies 10 – 15 ms and 15 – 21 ms. Following relevant previous work,^21,22^ we will refer to these subcomponents as the SN10 and Na, respectively. The other longer latency peaks (Pa, Nb) are dominantly cortical, but it is not clear at which peak (or at which latency) the switch from subcortical to cortical occurs. For instance, previous investigations of the SN10 using EEG and MEG^21–25^ interpret its likely sources as ranging from midbrain to thalamus to cortex. However, the P50 is more unambiguously generated by current dipole sources in the auditory cortices.^14^

An alternative approach to the traditional ABR and MLR, which additionally bypasses the limitations of relying on discrete stimuli but still maintains the interpretability of distinct response peaks (each with a characteristic polarity, magnitude, and latency), is to use continuous auditory stimuli (e.g., naturalistic speech) and the temporal response function (TRF) analysis framework.^26–29^ This newer but increasingly established methodology has been typically applied to cortical-only responses (e.g., P50 and later), but more recently has been applied to shorter-latency and higher-frequency responses including the ABR and MLR complexes.^30–34^ Remarkably, the TRF framework can produce both ABR/MLR-like TRF response profiles and longer-latency evoked-response-like TRF response profiles *from the same presentation of a single continuous auditory stimulus*, according to which stimulus feature is used in the TRF estimation process.

This result arises naturally since a TRF generated from a continuous stimulus requires the choice of one or more stimulus-feature regressors that are used to deconvolve the continuous response data. Indeed, different choices of stimulus-feature regressor give different TRFs, which can separately represent the neural processing of different stimulus features. For a typical cortical auditory TRF, the regressor is often chosen to be the overall envelope amplitude of the continuous stimulus, which generates TRFs with the familiar P50-N1-P2-like peak structure.^2^ For shorter-latency responses, however, it is critical to use a regressor that reflects the fine structure of the stimulus (i.e., that includes higher frequencies). For example, an auditory nerve model representation of the stimulus^35^ has already been extensively investigated and used as a regressor in this context.^33,34,36^ These EEG-only investigations consistently show a TRF peak with the same neural source as wave V, but using continuous stimuli such as naturalistic speech instead of averaging over click responses. Here, we replicate their findings, but greatly expand on them using simultaneous EEG and MEG recordings and subsequent neural source localization.

We recorded concurrent EEG and MEG responses while subjects listened to long duration stimuli. Most stimuli were 1-minute trials of continuous speech spoken by two talkers. The same subjects also listened to a 10-minute click train with varying interclick intervals. With both these categories of stimulus, we derived TRF response profiles for the overall speech stimulus (mixture of two speakers), and the clicks. Using the EEG+MEG functional data combined with each subject’s own structural MRI, a fine-grained source analysis was then performed using the EEG+MEG sensor-space TRFs to reveal the underlying neural current sources as a function of latency. These early-latency TRFs correspond to the ABR-MLR complex (Figure 1). Through a whole-brain threshold-free cluster enhancement (TFCE)^37^ permutation test, it can be established that 1) when the subjects were listening to continuous speech, the ABR-MLR TRF peaks were present at expected times, including wave V with a midbrain origin, followed by the SN10, Na, Pa, Nb, and P50, all originating from thalamic and cortical locations; 2) when listening to clicks, most, but not all, of the ABR-MLR TRF peaks were the same as when listening to speech, critically including the midbrain wave V, as well as some MLR morphology; and 3) there were noteworthy differences between speech TRFs and click TRFs at key times and brain regions, including within the SN10 and Na complex, as well as subsequent MLR peaks such as the Pa and Nb, at both thalamic and cortical sites. Most notably, we showed that the neural origin of the SN10 is cortical (with additional thalamic contributions) for auditory responses to speech, but not to clicks.

**Figure 1:**
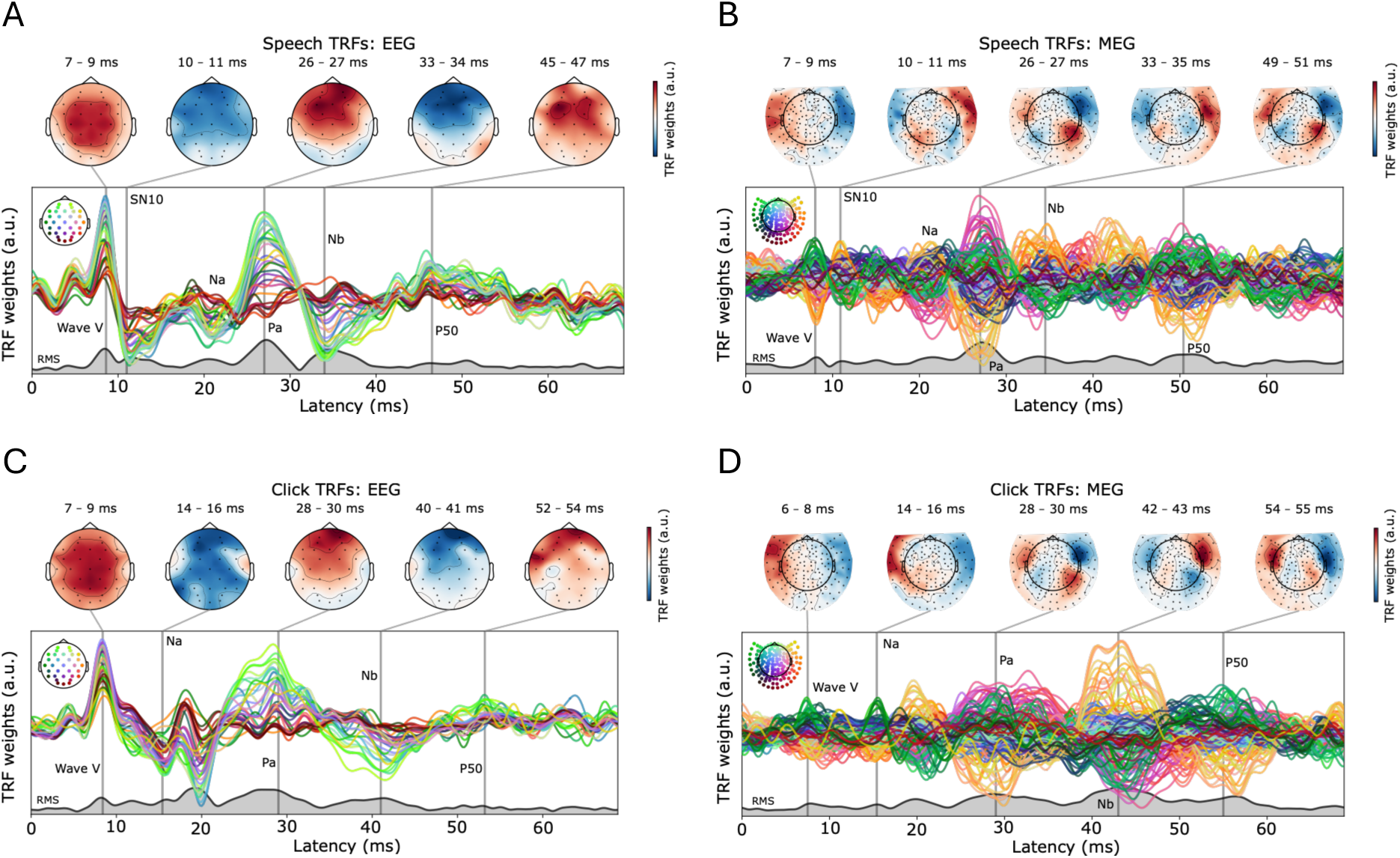
Sensor-space TRFs and select topographies for speech (top row: A and B) and click stimuli (bottom row: C and D). EEG-based TRFs on the left (A and C); MEG-based TRFs on the right (B and D). Prominent peaks can be identified as wave V (8-S ms), SN10/Na (11-15 ms), Pa (27-29 ms), Nb (41-43 ms), and P50 (46-55 ms). Note that the peak latencies vary across stimulus types (speech vs. clicks) and may not be identical to those of clinical ABR-MLRs.^17,20^

## Results

We recorded simultaneous EEG and MEG from 12 subjects (aged 18-29 years; 8 female) while they listened to 24 minutes of continuous speech from two competing speakers, as well as 10 minutes of click trains, all at a comfortable level of 70 dB SPL. The speech trials consisted of two competing speakers, where subjects were directed to listen only to one. The click train consisted of clicks with randomly varying interclick intervals of between 4 and 200 ms. This distribution of interclick intervals spans the range of typical inter-glottal pulse intervals in human speech, as well as longer interclick intervals more common in ABR or MLR research.

Among EEG researchers, it is more common to represent the ABR-MLR complex in sensor space (extracranially, e.g., in units of µV), whereas in the MEG literature it is more common to represent responses as currents in neural source space (per brain region, e.g., in units of nAm). As a nod to both, we first present sensor-space results and then present source-space results.

### TRF deconvolution in sensor space

To estimate any TRF, a stimulus representation/feature needs to be chosen as a regressor for reverse correlation with the neural response. To create these regressors, all audio stimuli (1-minute speech trials and 10-minute click trials) were processed with a well-established cochlear and auditory nerve model,^35,38^ as has now become common.^31,33,36^

Sensor-space auditory TRFs were obtained using the auditory nerve model as the regressor. Notably, these high sample rate TRFs were calculated with a full time-domain least-squares solution with cross validation, an approach more common in longer-latency cortical TRFs but otherwise underused for the earliest neural activity in the auditory system (see Discussion, particularly time-domain and frequency-domain approaches). Using this combination of approaches to deconvolution, while computationally expensive, ensures we obtain the greatest possible fidelity of responses. Prominent aspects of the ABR and MLR such as wave V, SN10, Pa, and P50, are clear in both EEG and MEG responses (Figure 1). Wave V of the ABR is most clearly visible in the EEG TRFs, for both speech and clicks, as expected,^32,33^ but is also visible even in the MEG TRFs, for both speech and clicks. The topographical properties of the wave V response are particularly evident, for EEG, showing a vertex-positive region centered at the top of the scalp, and, for MEG, showing a single medially-centered magnetic field dipole.

A key observation from Figure 1 is that the MEG topographies for later peak latencies, for example Pa (∼27 ms) and P50 (∼50 ms), show characteristic bilateral auditory cortex current source distributions, i.e., a prominent lateral magnetic field dipole over each hemisphere, with opposite polarity across the hemispheres.^39^ This same characteristic cortical topography also appears to emerge here surprisingly early, for the SN10 (∼11 ms), but only for the speech TRF. This manifests as an emerging cortical field configuration, indicated in the MEG topography by the opposite-polarity patches underneath the more frontal patches, and in the EEG topography as a slightly frontward-shifted center of mass of the field. This early indication of speech-specific auditory cortical activity will be further investigated below with source-space analysis.

### Sensor-space quantitative analysis

Cluster-based permutation tests showed a significant difference between speech and click TRFs, for both the EEG and MEG modalities. Figure 2 depicts a summary of these sensor-space TRFs with the first seven principal components for both modalities, with time ranges highlighted for clusters whose *p* values fall below 0.05. For EEG, two clusters discriminated between the speech and click responses: 1) from 30.6 to 36.4 ms, *p* = 0.00440, and 2) from 39.8 to 50.4 ms, *p* = 0.00049. For MEG, three clusters discriminated between the speech and click responses: 1) from 29 to 36 ms, *p* = 0.01954, 2) from 40.6 to 54.2 ms, *p* = 0.00195, and 3) from 47 to 63 ms, *p* = 0.02833.

**Figure 2:**
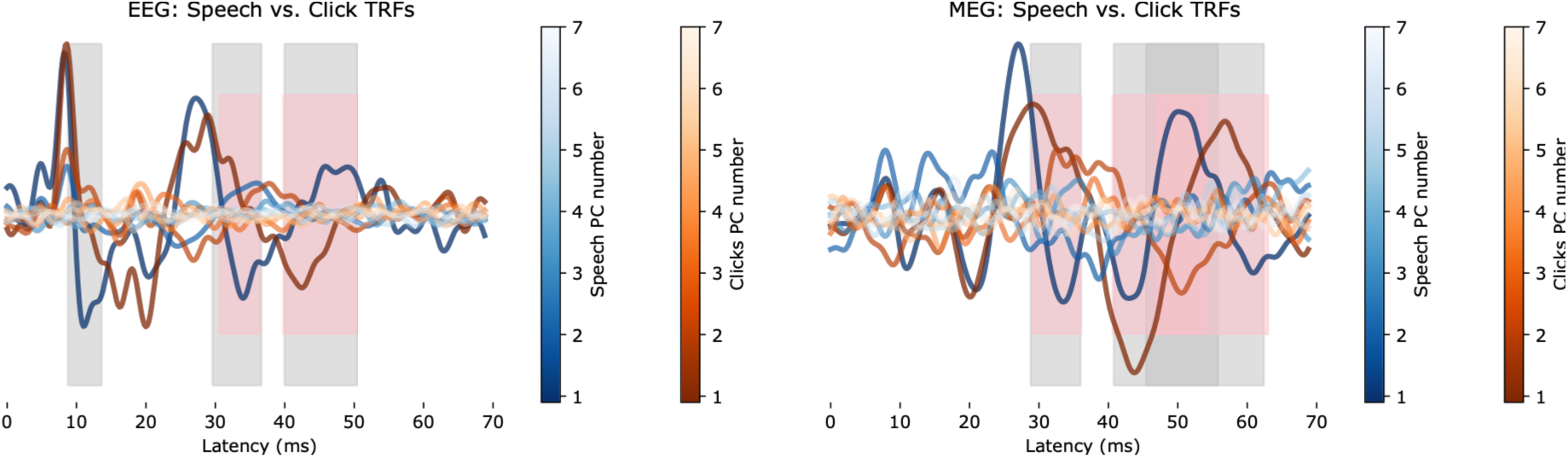
Sensor-space speech and click TRF principal components. Clusters comprising significant difference between speech and clicks are highlighted. Cluster times in pink are from time locking participant averages with reference to time zero, and clusters in silver are from time locking with reference to wave V latencies. The TRFs themselves, from which the PCs are derived, are averaged here with wave V latency time-locking. Increasingly lighter shades of blue and orange represent higher-order components of PCA, from number 1 through 7, for the speech and click TRFs, respectively.

This initial permutation test revealed no evidence that the ABR portion of the TRF (0 – 9 ms) contributes to any difference between speech and click TRFs, because no clusters with *p* < 0.05 were observed in that time range. However, there was a significant difference between the wave V latencies for speech (8.625 ms) and click (8.350 ms) TRFs via a simple paired-samples *t*-test (*t*(11) = 2.33, *p* = 0.040). In order to time-lock the subsequent MLR portions of the TRFs (10 – 60 ms) accurately to each other when testing for differences, we therefore instead time-locked the trial-averaged TRFs to their wave V latencies, for each participant and each stimulus type, for all further analysis. In other words, the MLR portions of the TRFs were time-aligned by first deconvolving with the auditory nerve model regressor representing the stimulus (analogously to a typical evoked response), and then as a finer-grained correction, realigning to their wave V, since the subsequent MLR portion of the response is itself initiated by the onset of wave V in the inferior colliculus.

The same spatiotemporal cluster permutation tests were then performed again after wave V-locking in sensor space, for both modalities and conditions. As expected, after this procedure there were still no significant differences found over the time period of the ABR (0 – 9 ms) with *p* < 0.05, indicating that the ABR does not contribute to any significant differences between our conditions. Critically, the significant differences between the speech and click TRFs during the MLR time period remained as such, plus one additional early cluster for the EEG responses. Also critically, the effect sizes of the original five clusters either remained the same (for the original two EEG clusters) or increased (for the three MEG clusters). Thus there are now, for EEG, three clusters that differentiate speech and click responses: 1) from 8.8 to 13.6 ms, *p* = 0.0186, 2) from 29.6 to 36.6 ms, *p* = 0.00440, and 3) from 40 to 50.4 ms, *p* = 0.00049; for MEG, three clusters that differentiate speech and click responses: 1) from 28.8 to 36 ms, *p* = 0.01856, 2) from 40.8 to 55.8 ms, *p* = 0.00147, and 3) from 45.4 to 62.4 ms, *p* = 0.02491. These clusters are summarized in Figure 3 and Table 1, and are also noted in the response principal components plotted in Figure 2; the TRFs from which the principal components in Figure 2 are derived are also averaged with respect to individual wave V times. Thus, the additional early cluster differentiating speech and clicks was specific to the EEG modality but fully encompasses the SN10 peak latency window. This, plus the MEG cortical topography at the SN10 time, motivated us to further pursue these significant differences between speech and click TRFs in source space.

**Figure 3:**
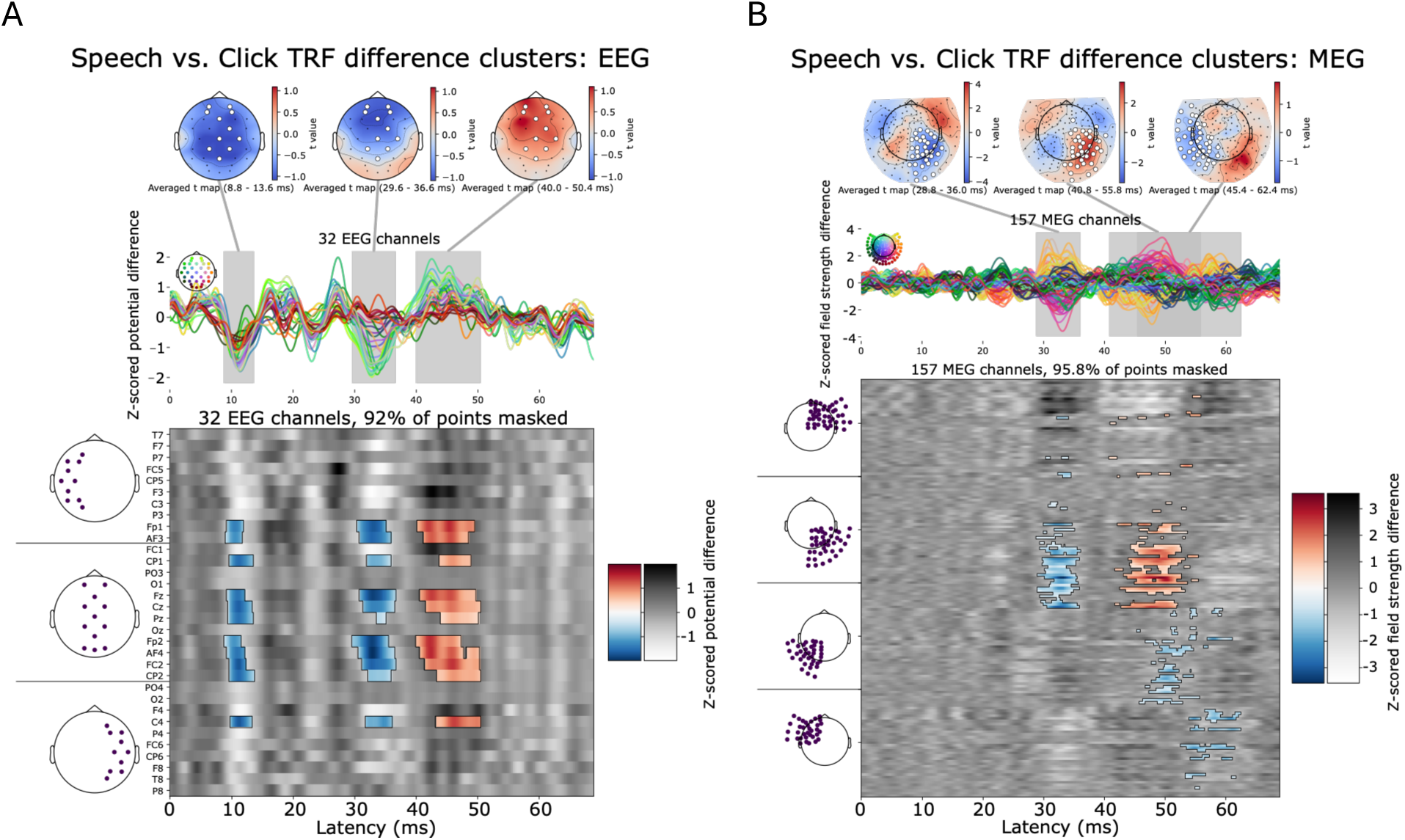
Sensor-space significant-difference clusters for EEG (A) and MEG (B). The difference between the sensor-space TRFs (listening to speech minus listening to clicks) are depicted in the butterfly time series plots and per-channel heatmap plots. Note that there is little difference evident during the ABR (including wave V) time period, as expected from the lack of significant-difference clusters between the speech and click TRFs at that time. The three topographies along the top correspond to the three main clusters of significant difference between the speech and click TRFs, also shown as silver background highlights along the time series of TRF differences. In the heatmaps of the sensor trace differences, cluster-member samples are shown in color from blue (negative) to red (positive), and non-cluster-member samples are shown in grayscale.

**Table 1:**
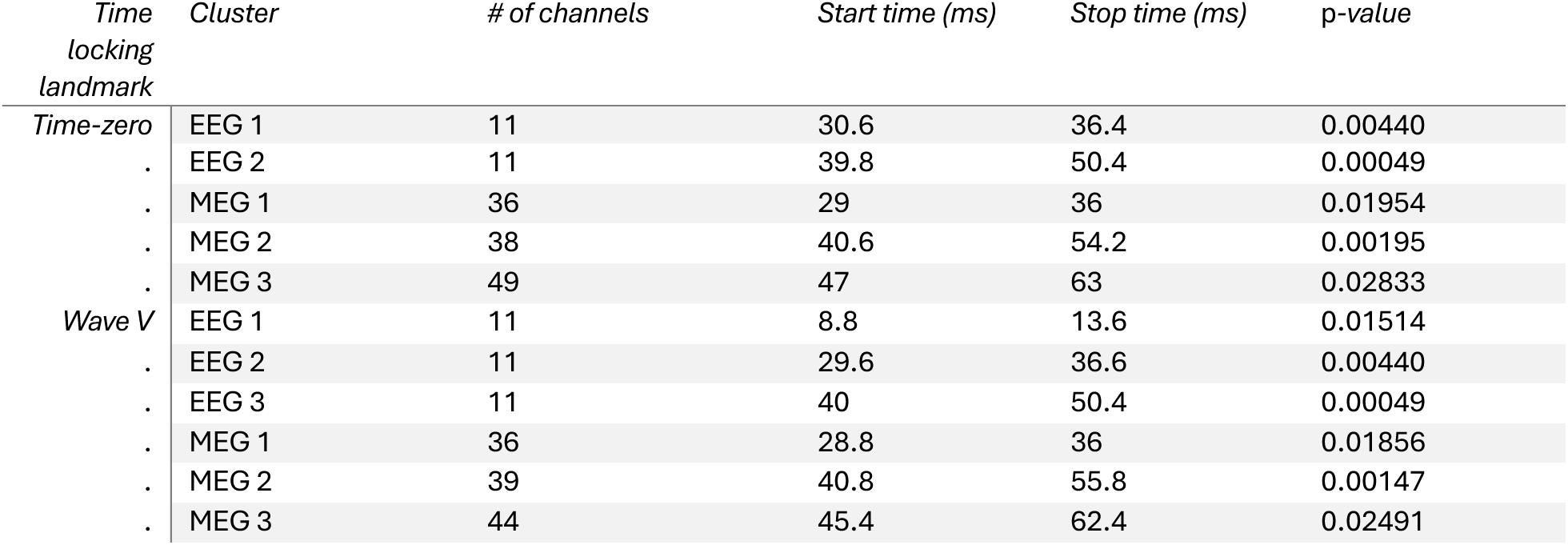
Sensor-space cluster statistics for differences between the speech TRF and the click TRF.

### Source-space analysis

In MEG sensor space, the TRFs showed that the SN10 has a bilateral topography, similar to the later peaks and consistent with bilateral cortical origin, but only for speech responses and not for clicks. To investigate the underlying neural current sources (including their anatomical orientation) of these and other responses, we undertook a fine-grained source analysis using all EEG and MEG sensors and a structural MRI from each individual subject. Source space volumes and forward models were constructed utilizing each subject’s brain segmentation, skin and skull surfaces, and sensor locations, and inverse modeling resulted in a minimum norm solution using MNE-Python.^40–42^ Both EEG and MEG recordings contribute to the source-space currents obtained, but not necessarily equally, given their physical weighting from Maxwell’s equations and the anatomical models, resulting in EEG having greater sensitivity to deep sources. Broadly, EEG is expected to contribute more towards the localization of deeper sources, such as the midbrain and thalamus, while MEG is expected to contribute more to the ability to differentiate cortical sources, including any cortical hemispheric asymmetry.

We tested for significant differences between each TRF and 0, and for significant differences between the two TRFs. We again first synchronized the wave V latencies for each TRF, time-locking the source-space time series per participant and per condition to the wave V times before further source-space analysis. After transforming the sensor-space TRFs into the volume source spaces (maintaining the full vector-valued, three-dimensional neural current source solutions), we subjected the TRFs to a whole-brain TFCE cluster analysis^37^ to test for differences between each condition and zero (one-sample tests) and between conditions (paired-samples test). Hotelling’s T-squared statistic was used in order to simultaneously test all three Cartesian dimensions of the vector-valued solution.^43^ We note that, in the case of the paired-samples test, if a significant difference between the two TRFs is found for source space currents, the cluster at the given time and location indicates that: a) TRF current magnitudes of those two TRFs were different at that time and location, or b) TRF current orientations of those two TRFs were in different directions at that time and location, or c) some combination of both.

### Significant clusters in source space

For the one-sample tests for significant activation, there were multiple clusters showing a significant difference between both TRFs and zero. While *p*-values for individual clusters from the TFCE test are already corrected, for further analysis we employed a more conservative threshold by only considering corrected *p*-values below 0.02 (the effect of which is to omit clusters with small number of vertices or with time windows ≤ 1ms). Utilizing this threshold, there were four partially overlapping clusters contributing to the speech TRF’s difference from zero, and one long cluster contributing to the click TRF’s difference from zero. The temporal extent of the speech TRF clusters was continuous from 1 to 44 ms (see Table S3), and for the click TRF cluster was continuous from 0 to 64 ms (Table S4).

Figure 4 shows some prominent spatiotemporal locations where the two TRFs are significantly different from zero (overall time periods of clusters contributing to the significant differences between the two TRFs and zero are depicted in Figure 5). In these plots, only vertices which are part of the cluster are shown on the brain. Figure 4A shows the ABR wave V in its expected midbrain location (see also Figures S1 and S2), while the SN10 negativity is shown in 4B. Already at 11 ms, the TRF current source has spread to auditory cortex as revealed here. The Na negativity shown in 4C confirms a clear cortical origin by 21 ms for the speech TRF. Figure 4D shows the P50 component from the click TRF, with a bilateral cortical origin.

**Figure 4:**
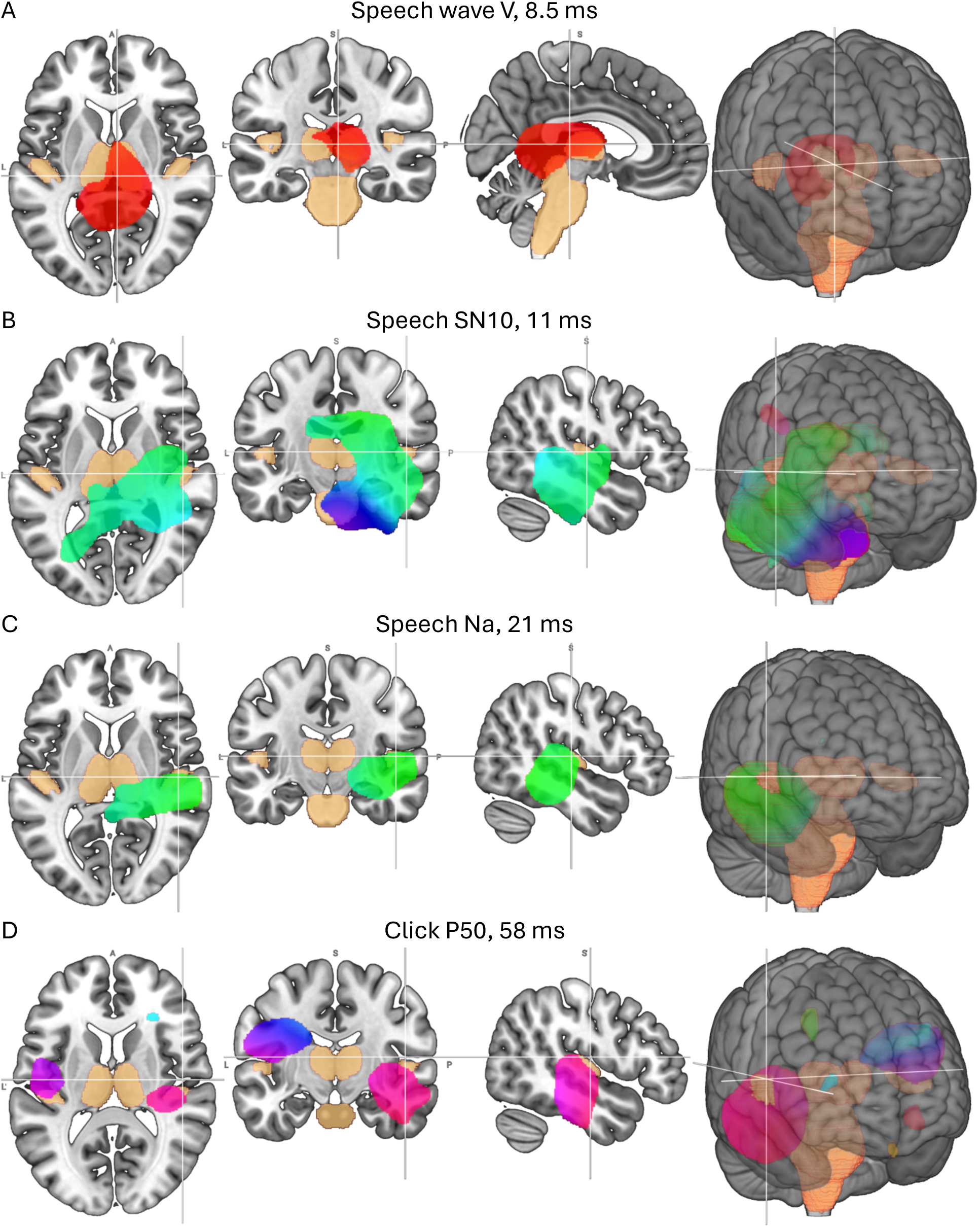
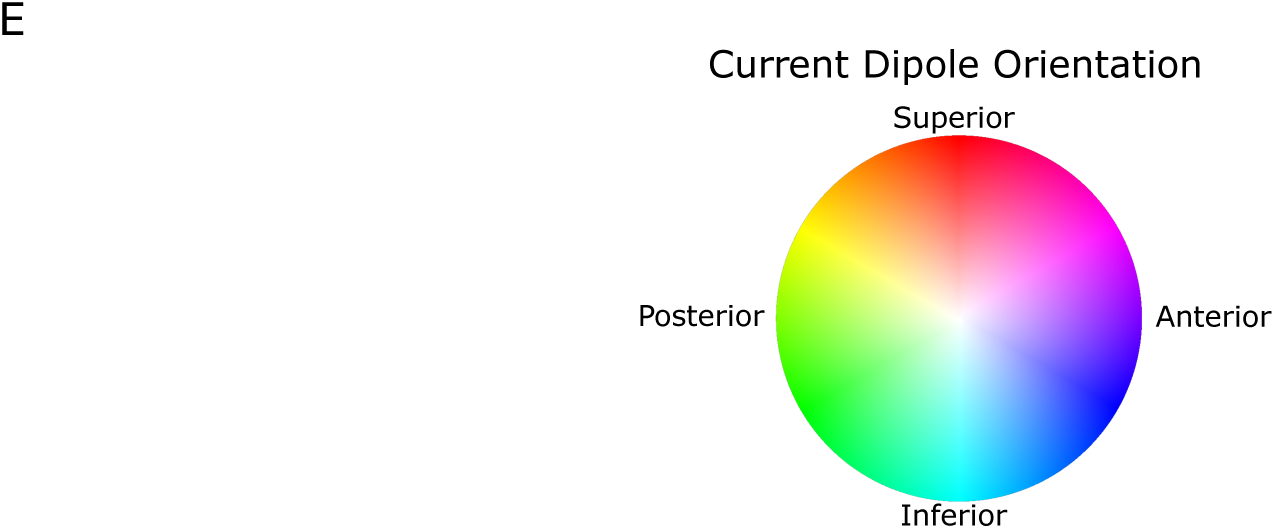
Several spatiotemporal clusters for which the speech and click TRFs are different from zero. While not representative of all times and places contributing to the TRFs’ significance, these clusters demonstrate robust TRF peaks that are different from zero, and also for some (A, C, and D), for which the speech and click TRFs do not significantly differ from each other (cf. Figures 5 – 7). MRI slice images use the neurological convention (right hemisphere shown on the right) throughout this work. MRI 3d renders, e.g. the rightmost images of each subfigure, are typically viewed from the front looking posteriorly, however, thus these are technically in radiological convention (right hemisphere shown on the left). (A) Wave V for the speech TRF, at 8.5 ms. Magnitude is centered in the midbrain around the inferior colliculus as expected, with current orientation straight up toward the vertex. B) SN10, also for the speech TRF, at 11 ms. Magnitude has quickly spread to the auditory cortex while also remaining in the thalamus and midbrain regions. This early cortical source contributes to the speech TRF’s significant difference from zero, and also to its difference from the click TRF (Figure 7). C) Na, also for the speech TRF, at 21 ms. By this time, the TRF source is clearly shown to be auditory cortex, with current orientation inferior and posterior, consistent with the vertex-negative character of this typical part of the MLR. D) Late P50 for the click TRF. This shows bilateral auditory cortex activation, with orientation superior and anterior, consistent with the vertex-positive character of this part of the MLR and also with the orientation of Heschl’s gyrus. Anatomical landmarks derived from the Harvard-Oxford volumetric Cortical Structural Atlas (RRID: SCR_001476) as distributed with the FSL software^44,45^ are shown in gold: brainstem, left and right thalamus, and left and right primary auditory cortex. Current vector directions are projected into two dimensions: superior-inferior and anterior-posterior, mapped onto the circular colormap shown in (E). Current magnitude is represented by the percentage of total source space vertices plotted. Thus, a cluster of larger magnitude also appears larger in its extent, while a cluster of smaller magnitude appears smaller in extent. Additional spatiotemporal clusters for which click TRFs are significantly different from zero are shown in Figure S2.

**Figure 5:**
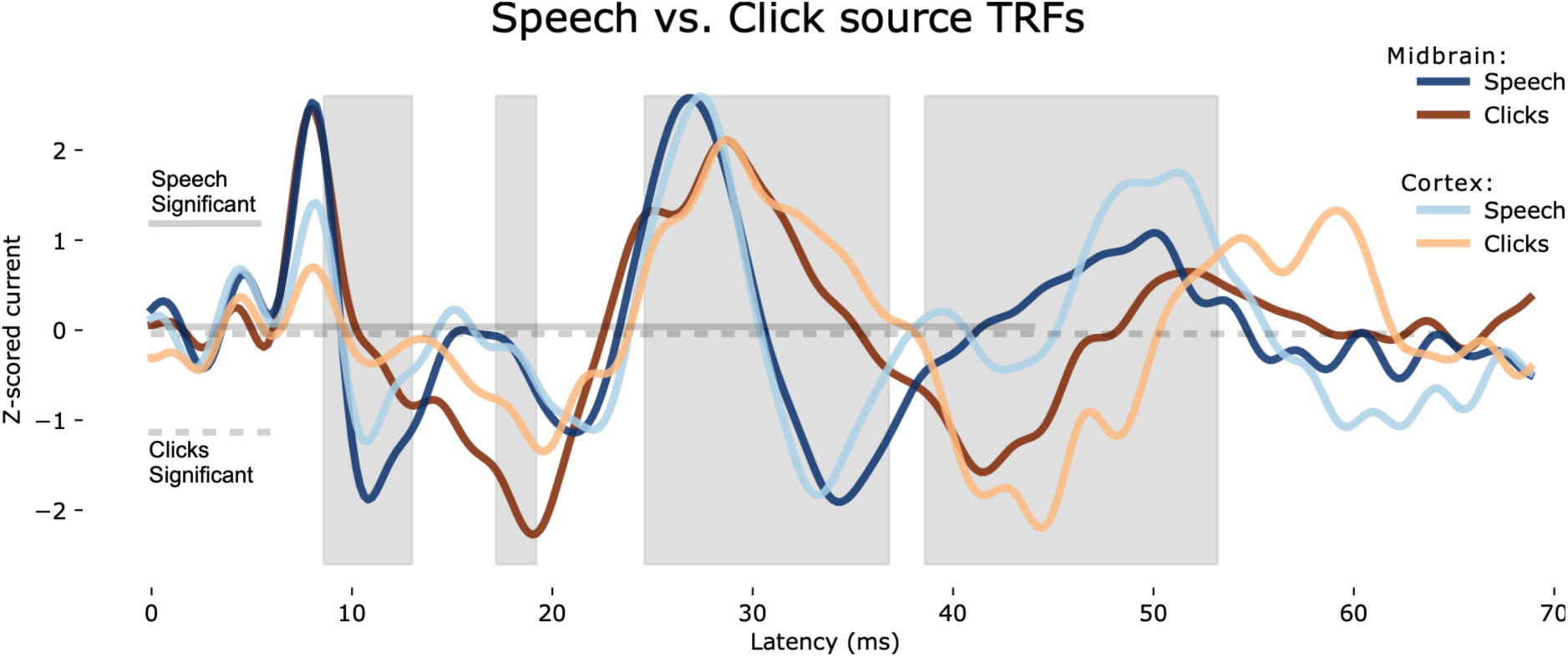
Source-space time courses for midbrain and primary auditory cortical regions. Clusters contributing to significant difference between speech and click TRFs (from Table 2) are highlighted in silver. The ABR time period (∼0 – S ms) does not significantly differentiate these two TRFs in source space (as also the case in sensor space), subsequent clusters of difference demarcate similar MLR structures as in the sensor space analysis: SN10, Na, and Nb peaks all contain differences between speech and click TRFs, with lesser but still significant differences at the Pa and P50 times. The solid and dashed lines indicate the times that the speech and click TRFs are independently different from zero, respectively.

**Table 2:** Source-space cluster statistics for differences between the speech TRF and the click TRF. Clusters are sorted by p-value ascending, and all clusters with p < 0.05 are reported in this table. Though all TFCE-based clusters are significant with multiple comparisons correction having been applied, only clusters with p < 0.02 were considered for further analysis and visualization (e.g., Figures 6 and 7). While arbitrary, this avoids over-interpreting significant clusters with a small number of vertices, or windows of duration ≤ 1ms.

| *# of source vertices* | *Start time (ms)* | *Stop time (ms)* | p-*value* |
|---|---|---|---|
| *1499* | 24.6 | 36.8 | 0.00147 |
| *457* | 8.6 | 13 | 0.00269 |
| *1191* | 38.6 | 53.2 | 0.00952 |
| *54* | 17.2 | 19.2 | 0.01148 |
| *2* | 57.4 | 57.8 | 0.03443 |
| *1* | 53.4 | 53.8 | 0.03712 |
| *10* | 29.8 | 30.8 | 0.03956 |
| *1* | 53.8 | 54 | 0.04053 |
| *1* | 52.8 | 53 | 0.04151 |
| *1* | 55.4 | 55.6 | 0.04322 |
| *10* | 53.8 | 54.4 | 0.04371 |
| *5* | 29 | 29.6 | 0.04493 |
| *6* | 48.4 | 49 | 0.04640 |
| *1* | 53.4 | 53.8 | 0.04640 |
| *1* | 42.6 | 42.8 | 0.04640 |
| *1* | 53.4 | 53.6 | 0.04640 |
| *1* | 53.4 | 53.6 | 0.04884 |
| *1* | 54.8 | 55 | 0.04884 |
| *1* | 53.4 | 53.6 | 0.04933 |
| *1* | 29.4 | 29.6 | 0.04933 |
| *1* | 53.6 | 53.8 | 0.04957 |
| *1* | 54.6 | 54.8 | 0.04957 |
| *1* | 53 | 53.4 | 0.04957 |
| *1* | 55.6 | 55.8 | 0.04982 |

For the statistical comparison between the speech TRF and the click TRF (formulated as speech minus click), there was a significant difference between the TRFs, represented by a total of 24 spatiotemporal clusters (Table 2). With further analysis again employing the more conservative threshold of corrected *p*-values below 0.02 (the effect of which is to omit clusters with small number of vertices or with time windows ≤ 1.2 ms), however, four clusters still remained. An accounting of each cluster follows below, in reverse temporal order, from the well-understood P50 back down to the enigmatic SN10. As seen below, significant differences may be due to instantaneous differences in magnitude, shifts in peak latency, or combinations of both.

Summary source-space time series and cluster timing is depicted in Figure 5. Whereas the cluster tests depicted in Figure 4 (and other figures below) show the exact locations of source-space clusters at specific times, source-space summary time courses may give some visual intuition for the transformation from sensor to source space. In this figure we have extracted the source-space time-courses^46^ both from primary auditory cortex and from midbrain and thalamus in order to 1) visualize the time courses of the speech and click TRFs in source space, 2) visualize the timing of clusters showing significant difference between the speech and click TRFs (analogous to Figure 3 for sensor-space clusters), and 3) show how time courses from medial and cortical sources differ.

In the following sections we will present the significant differences between the speech and click TRFs in reverse temporal order, i.e., from later to earlier.

### Late MLR thalamocortical and cortical processing

*Nb and P50 cluster, 38.6 to 53.2 ms* (1191 source vertices, *p* = 0.01): This cluster of significant difference between speech and click processing comprises activity in both thalamus and bilateral auditory cortex. There are two categorical substages of this cluster, the P50 stage centered on 49 ms (Figure 6A) and the late Nb stage at 43 ms (Figure 6B). At 49 ms, the speech TRF has greater magnitude than the click TRF, and with a superior-anterior current orientation; the difference is therefore largely due to the presence of the speech TRF P50. The click TRF P50 is later (56 – 59 ms), though there is no significant difference between the TRFs at that time. At 43 ms (Figure 6B), the difference is largely due to the presence of the click TRF Nb. The magnitude of the speech TRF at this time is near zero, and the orientation of the click TRF is vertex-negative. The significant difference (speech TRF minus click TRF) is vertex-positive, indicating that the click TRF has greater activity, as its orientation is inferior-posterior. In this light, the late Nb stage can be identified as click-generated Nb.

**Figure 6:**
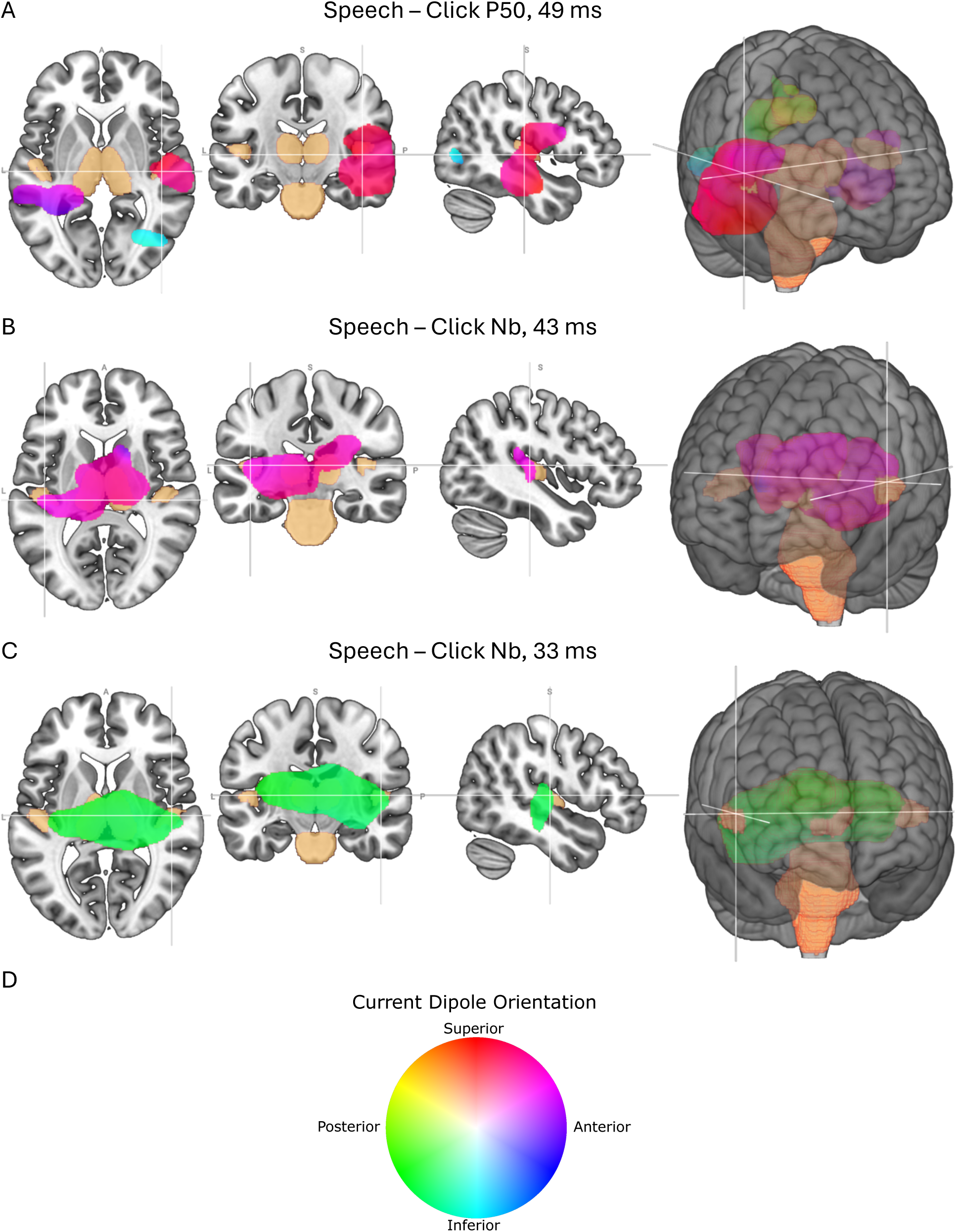
Later clusters of significant difference between speech and click TRFs, in order of decreasing latency, using the same conventions as Figure 4. Slice and render orientations as in Figure 4. A) P50 stage at t = 49 ms, where the response difference is dominated by the speech TRF P50, with superior-anterior current orientation, whereas the click TRF P50 has not yet begun; B) Late Nb stage at 43 ms, where the response difference is dominated by the click TRF Nb, with vertex-negative current orientation, and an absence of a speech TRF late Nb; C) Early Nb stage at t = 33 ms, where the response difference is dominated by the speech TRF Nb, with vertex-negative current orientation, and an absence of a click TRF Nb until later. Current vector directions are mapped onto the circular colormap shown in (D).

*Pa and Nb cluster, 24.6 to 36.8 ms* (1499 source vertices, *p* = 0.001): This is the largest cluster in both duration and magnitude, comprising much of thalamus and bilateral auditory cortex. As shown in Figures 5 and 6C, centered on 33 ms, the speech TRF has greater magnitude, with vertex negative polarity, while the magnitude of the click TRF is near zero. The difference is vertex-negative, indicating that the speech TRF has greater activity. This vertex-negative current dipole appears to be the Nb of the speech TRF, which would put it earlier than typically found in the clinical (click-train generated) MLR literature. In this light, the early Nb stage can be identified as the speech-generated Nb, the latency of which is earlier than for discrete (non-speech) sounds.

*Late Na cluster*, 17.2 to 19.2 ms (54 source vertices, *p* = 0.011): A cluster dominated by activity in the thalamus, as seen in Figure 7A. The click TRF has greater magnitude and the significant difference is vertex-positive, whereas both TRFs are vertex-negative. This indicates that the Na is generated by thalamic activity present in both TRFs, but greater for the click TRF.

**Figure 7:**
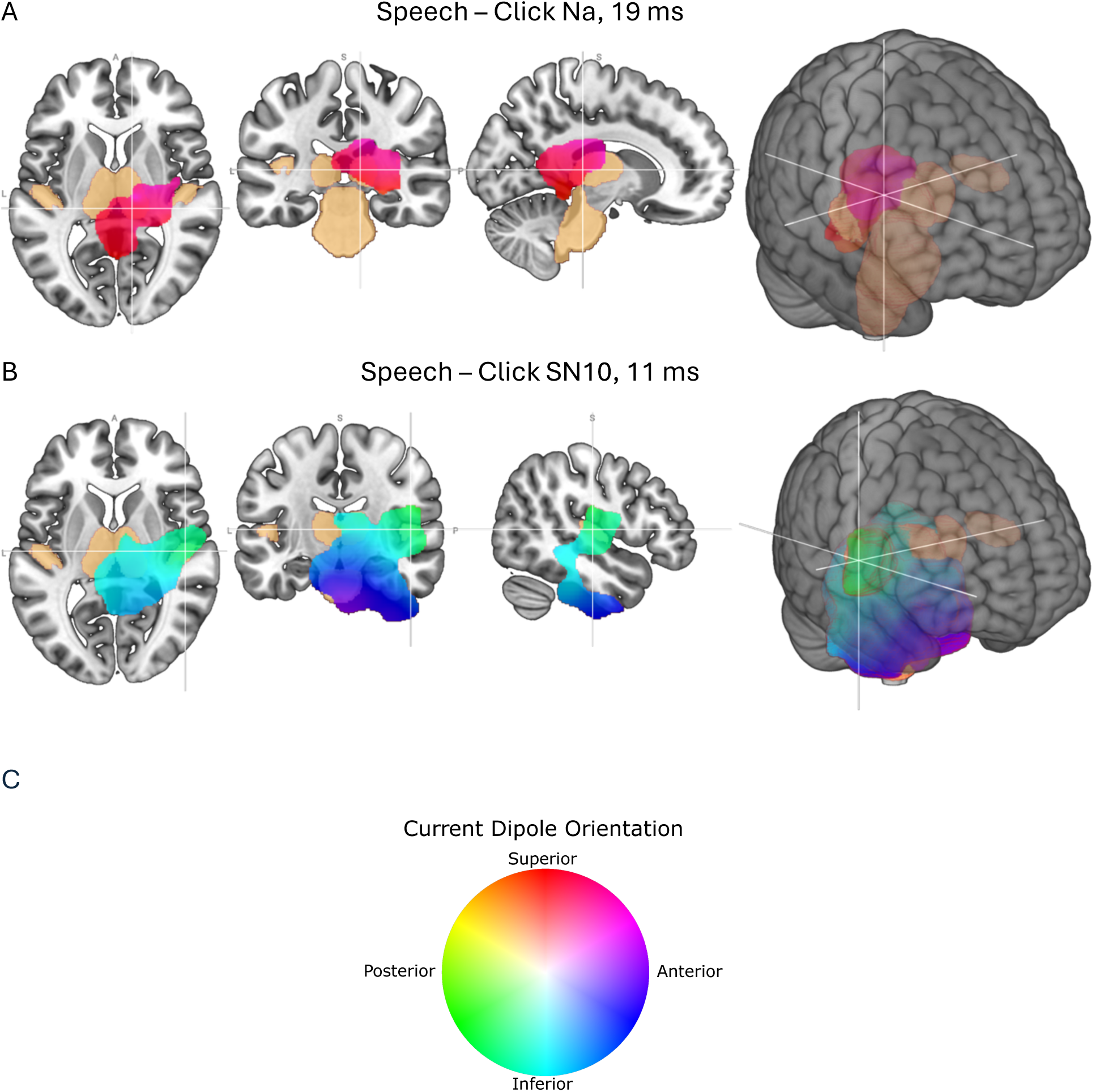
Earlier clusters of significant difference between speech and click TRFs, in order of decreasing latency, using the same conventions as Figure 4. Slice and render orientations as in Figure 4. A) Late Na stage at 19 ms, where the response difference is due to a larger click TRF relative to the speech TRF, both with vertex-negative current orientation; B) SN10 stage at 11 ms, where the response difference is due to the presence of a speech TRF SN10 originating dominantly from thalamus and auditory cortex, with vertex-negative current orientation, and no such contribution from the click TRF. This indicates that this very early thalamocortical activity is present for speech but not clicks. Current vector directions are mapped onto the circular colormap shown in (C).

*SN10 cluster, 8.6 to 13 ms* (457 source vertices, *p* = 0.003): Despite being so early, this cluster is dominated by thalamus, auditory cortex, and other intermediate regions (Figure 7B). This cluster is oriented in the vertex negative direction, and the speech TRF has greater magnitude than the click TRF, so the speech SN10’s orientation is the same as the significant difference. Note that the distribution of SN10 peak magnitudes across individuals for both speech and click TRFs, as illustrated in Figure S10, also shows a significant difference across stimuli, via a simple paired-samples t-test (*t*(11) = 4.51, *p* = 0.00088). Part of this cluster is located inferior to the thalamus and oriented anteriorly and inferiorly, similar to the analogous cluster for the speech TRF by itself (Figure 4B); while it is possible this location of activity represents fast efferent current moving back down the auditory pathway, it is also important that the magnitude at this location is markedly less than the dominant thalamocortical vertices of the cluster, as shown in Figure S7 with its more conservative threshold. In short, this very early (∼11 ms) activity is thalamocortical (both thalamic and cortical) and is substantially present during speech processing and not clicks.

## Discussion

The use of continuous speech as a stimulus, with an auditory nerve model to generate TRFs, in concert with simultaneous EEG and MEG recordings, has given us unprecedented access to the neural origins of early latency (0 – 60 ms) stages of subcortical and cortical auditory processing. Most notably, the SN10 processing stage provides compelling evidence for the earliest cortical response (11 ms) yet seen noninvasively. Critically, this response, originating from both thalamus and auditory cortex, is present only for the speech stimulus but not the click train stimulus.

This speech-specific processing stage might result from several factors, including task-based plasticity, speech processing experience over the participant’s lifetime, or evolution-based speech sensitivity in humans. This in turn raises the question of what statistical properties of the speech sounds are actually tracked by this mechanism. Candidate properties include the particular autocorrelation function of the stimulus (interclick or inter-glottal pulse interval distributions being an operationalization of this property; see Figure S9), pitch fluctuations (prosody trajectories of speech sounds or similar non-speech sounds as an example), timbre (spectral distributions), or of course top-down aspects such as the semantic and pragmatic information that a coherent speech utterance contains. Most of these possibilities invite empirical approaches to address this question but are not testable in the current study. Future efforts could systematically test such possibilities by varying stimulus parameters of the click train, either as further discussed below, or something more complex, along the above dimensions, including “naturalness”. One could also vary parameters of the speech stimulus, altering the natural, synthesized speech of the type used here to become less naturalistic and more click-like; in this approach, the participant would always hear intelligible speech, while multiple auditory parameters could alter its statistical structure. Such experimental manipulations would then be able to disentangle which stimulus or experimental design properties lead to earlier auditory cortical contributions.

While this is the first source analysis study of early-latency auditory responses using continuous speech (and the TRF paradigm), there are many varied reports in the literature investigating the latencies and sources of these response components using traditional evoked paradigms. Using MEG alone,^24^ sources of the click-evoked SN10 were localized, placing it more medially than subsequent MLR stages but more lateral than thalamus. Other MEG source analysis placed the earliest cortical component of the click-evoked ABR-MLR complex at a late Na time of ∼17–19 ms.^23,47^ More recently, also using MEG,^22^ the click-evoked SN10 was robustly recorded, but the equivalent current dipole location was found in the midbrain, in contrast to Na and later peaks which were well-modeled by auditory cortical sources. These earlier source analyses of the click-evoked SN10 thus paint a somewhat inconsistent picture, particularly when compared to the earlier wave V, near-universally regarded as having its source as the inferior colliculus, and later processing stages (Pa and subsequent waves, which are more consistently cortical). Thus our result that the SN10 appears to be speech-specific provides some clarification and context to the literature; even apart from specific neural sources, it is noteworthy that the morphological differences between the speech and click TRFs are robust around this timepoint.

Related work has also utilized electrocorticography (ECoG) in awake humans to record auditory TRFs to natural speech directly from neurons in and around the auditory cortex.^48^ This work uncovered important facts about auditory responses to continuous speech, including the existence of both “onset” and “sustained” response profiles in neurons throughout the superior temporal gyrus. While these onset-profile neurons were significantly earlier from their sustained-profile counterparts as quantified by TRF latency, their relationship to the present work has two important aspects: they were not characterized by latencies as early as the SN10 reported here, instead clustering around the more common P50 time, and also, while the experiment concentrated on results from speech stimuli, these neurons responded equivalently to non-speech stimuli. Other relevant and more recent work in the mouse auditory system has shown faster (than otherwise expected) latencies in the cortex that arrive via alternate pathways, specifically a non-lemniscal pathway projecting to layer 6, rather than layer 4, of both primary and secondary auditory cortex.^49^ There, the subcortical structures responsible for information transmission through this non-lemniscal pathway also receive large numbers of efferent, top-down projections, indicating that behaviorally and otherwise ethologically relevant modulation may be relevant in this network. Considering the present speech-specific evidence, and its relationship to both the present and previous click-stimulus-related evidence, this non-lemniscal projection to layer 6 of auditory cortex might underlie a physiological mechanism of speech sensitivity in the human auditory brain. While the short-latency pathway in mouse was observed for a wide range of stimuli including tones and clicks, it was also seen that frequency tuning among units in layer 6 in this pathway is broader than that of the layer 4 counterparts. This may be consistent with the faster latency seen in the current data.

If this study’s speech responses are different than the click responses because of reasons along the lines of the above, this could also explain the significant differences we see between the speech and click responses subsequent to the SN10. The thalamocortical loops responsible for the MLR after this stage may be modulated by the pathway and latency differences caused by the speech nature of the stimulus, for instance allowing the Nb to occur approximately 10 ms earlier and with a different morphology for speech responses than click responses. However, the MLR literature suggests there is also variability of the Nb and P50 latencies as a function of stimulus rate, and the ABR literature certainly shows that wave V latency varies with click rate, and even sound level.^18,19^ In the present work, we record wave Vs that are slightly earlier for clicks (8.35 ms) than for speech (8.625), while at least two of the later MLR peaks appear both different in morphology and possibly later for clicks than speech. While the interclick intervals used for the click trains here include the range of human speech fundamental frequencies (approximately 80 – 200 Hz), the median frequency of interclick interval for the click trains here was only 34 Hz, whereas the comparable number for speech is approximately 163 Hz, including both male and female speakers. With this in mind, our slight difference in wave V latencies is in line with previous literature, as wave V latency tends to be a monotonic positive function with stimulus rate.^16,18,19^ Interestingly, the Nb and P50 have a nonmonotonic relationship with stimulus rate, with low rates (∼5 Hz) eliciting shorter latencies (41 and 52 ms respectively), medium rates lengthening the latency (∼39 Hz leading to 42 and 59 ms), and fast rates again decreasing the latency (∼100 Hz leading to 39 and 50 ms).^17,20^ This is also consistent with the current dataset; from the stimulus rate perspective, we only employed medium and high rates, but the MLR latencies are longer in our data for clicks (medium rate) than for speech (high rate), which may be partly responsible for the overall significant difference between these two TRFs. For speech, however, the Nb latency at ∼33 ms is substantially earlier than what is common in the MLR literature for clicks, and, most interestingly, the SN10 effects seen here are categorically different than those typical in click-evoked MLR investigations, both of which contribute to the overall significant difference.

Other work relevant to latency measures in human primary auditory cortex comes from ECoG. Indeed the fastest latencies recorded invasively are comparable to those seen here, at 11.9 ms^50^ in one and 10 ms^51^ in another. In both of these cases, electrodes along the most medial portion of Heschl’s gyrus yield the earliest responses, which is also consistent with the findings here of a thalamocortical source that, while overlapping the Heschl’s gyrus ROI, is also not as lateral in its centroid as, e.g., auditory P50 responses seen here and elsewhere. These early responses measured in primary auditory cortex are part of a large distribution of latencies, ranging from 10 to 100 ms. Both of these invasive studies employed click stimuli to evoke their earliest responses, which were not observed in this noninvasive study. Interestingly, one of these studies^51^ also employed single-syllable speech stimuli, but the corresponding evoked responses showed longer latencies (26 ms) rather than the early latencies seen here for continuous speech.

Our ability to record robust ABR-MLR responses as TRFs in a continuous speech paradigm builds on recent important work^10,11,30,32,33,52^ in several ways. Here we deconvolve TRFs without regularization, similar to approaches that use frequency-domain computations for speed.^33,34^ While the frequency-domain approximation of least-squares regression in TRF paradigms is highly useful, it is crucially only valid for stimulus representations with stationary statistics,^53^ whereas the more general time-domain form is necessary when stationarity cannot be assumed. The difference here regards the stimulus autocorrelation representation: For stationary statistics, this can be approximated in the Fourier domain with the simple spectrum of the stimulus regressor, whereas the full autocorrelation matrix (generally the Gram matrix of the canonical least-squares solution, Equation 1) over the entire time course of the TRF is needed for the non-stationary case. Thus, while the cross-correlation portion of the regression can always equivalently be performed as the cross spectrum in the Fourier domain, conversion back to the time domain in order to invert the autocorrelation matrix is necessary in the general case of long duration speech stimuli.

Along these lines, here we use the full time-domain least-squares solution, which is similar to recent work that also uses time-domain deconvolution for fast auditory TRFs, but which also typically employs either regularization or full cross correlation.^10,11,54,55^ While regularization or cross correlation can reduce overfitting, they also introduce an admixture of the stimulus autocorrelation function into the TRFs, to at least some extent, by design. In the general case of regularized regression, there is a danger of some TRF peaks occurring, or peak latencies being modified, due to the influence of structure in the stimulus autocorrelation function; this would be observed as artifactual peaks in the TRF that are actually due to acoustic features at shifted latencies (by one or more periods of the autocorrelation). The linear least-squares solution avoids these confounds by using the full, non-regularized stimulus regressor autocorrelation matrix.

As the click train utilized here employed randomly varying interclick intervals, while the speech stimuli were naturalistic, the temporal features including the autocorrelation functions of these two stimuli (and their respective auditory nerve model regressors) are different (Figure S9). While the click train regressor has little autocorrelation at any lag, the speech regressors have characteristic, broad peaks in their autocorrelation functions (approximately 5 ms and 10 ms) corresponding to the average fundamental frequencies of the female and male speakers. While this categorical difference in temporal structure could contribute to the differences in the resulting TRFs, the method of using linear least-squares regression without regularization ensures that temporal features in responses due to autocorrelation in the regressors are minimally present, because of the division by the autocorrelation of the regressor in Equation 1.

It is possible that a click train constructed to more accurately approximate the temporal and spectral attributes of speech (such as autocorrelation as a result of interclick intervals mimicking the slow and steady change of glottal pulse frequencies) may yield a different TRF. However this possibility would not preclude the explanation offered here of the cortical SN10 being a top-down effect of speech. Indeed, if a click train stimulus was constructed to closely mimic natural speech, the brain may indeed process it as if it were speech. Future research on this specific aspect of the stimuli could yield fruitful comparisons.

The early cortical component of the speech TRF here also leads to the question of what other stimulus types with ethological relevance may generate a comparable response. Natural sound textures and music are among stimuli that should be further tested along these lines, although human speech has the advantage of being composed of relatively punctate subparts, in its glottal pulses, where the waveforms of natural textures and musical sounds tend to be smoother. Recent evidence, however, shows that speech and musical sounds are largely similar in their subcortical and faster-latency TRF components, while their cortical and longer-latency responses diverge.^34^

As this is the first study to use time-domain least-squares regression at a high sampling rate, as well as incorporating neural source localization using both EEG and MEG, our data provide a level of detail on these phenomena not yet seen. In addition to robustly recording the sensor-space ABR for both EEG and MEG, and with comparable clarity to previous ABR literature, we are also able to differentiate between TRFs to different stimulus types with a fine-grained source analysis, and, notably, provide compelling evidence for the earliest noninvasive cortical response (11 ms) yet seen in humans.

## Resource Availability

### Lead Contact

Karl D. Lerud

## Materials Availability

The custom code to conduct these analyses is archived at https://doi.org/10.5281/zenodo.23813431 and is also available at https://github.com/lerud/eegAndMegAnalysis/tree/releases

## Data Availability

The data collected for the analyses in this work are archived at https://doi.org/10.5281/zenodo.23773280

## Acknowledgements

This work was supported by the National Institute on Deafness and Other Communication Disorders (NIDCD) grant R01-DC019394 and training grant T32-DC00046 (to CF and VC). The authors also acknowledge the University of Maryland supercomputing resources (http://hpcc.umd.edu) made available for conducting the research reported in this manuscript.

## Author contributions

Conceptualization: K.D.L., J.Z.S.; Data collection and curation: K.D.L., C.F.; Funding acquisition: J.Z.S.; Investigation: K.D.L.; Methodology: K.D.L., C.F., V.C., S.A., J.Z.S.; Resources: S.A., J.Z.S.; Software: K.D.L., C.F., V.C.; Supervision: J.Z.S.; Validation: K.D.L., C.F., V.C.; Visualization: K.D.L.; Writing: K.D.L.; Writing – Review C Editing: K.D.L, C.F., V.C., S.A., J.Z.S.

## STAR*Methods

### Key resources table

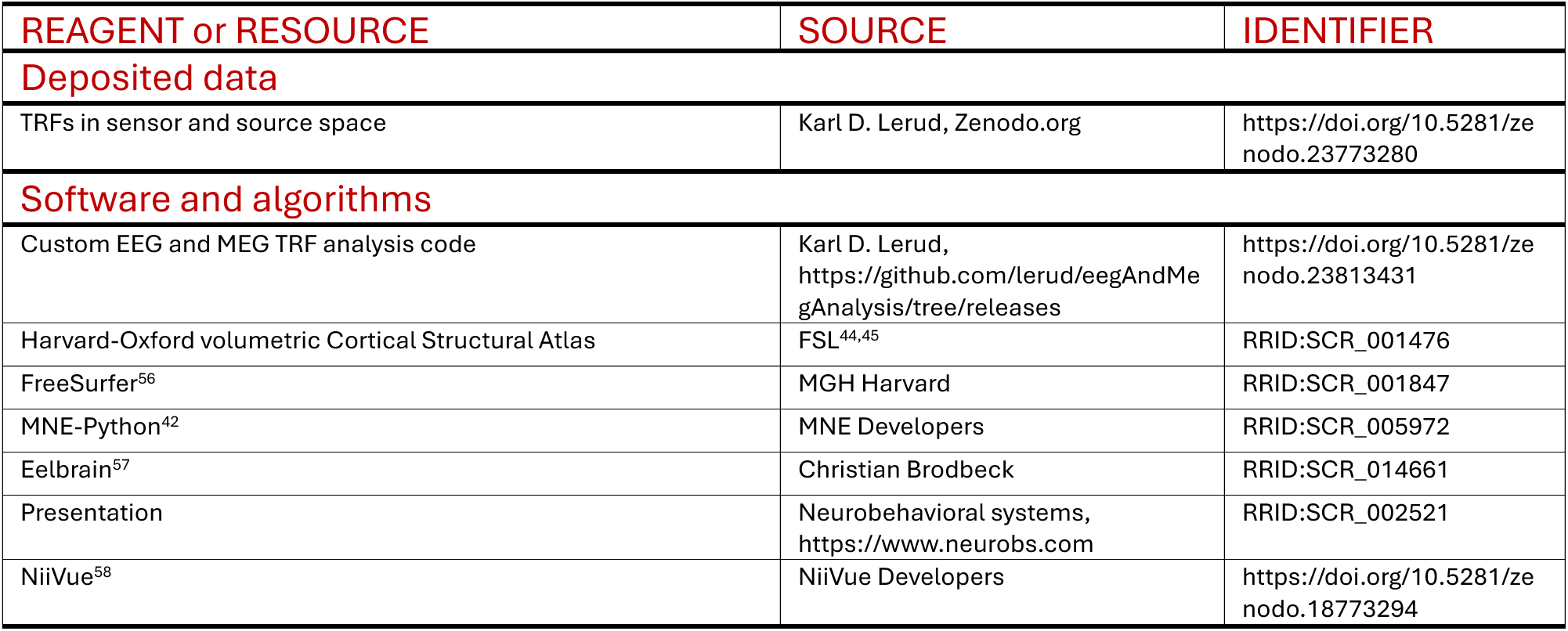

| REAGENT or RESOURCE | SOURCE | IDENTIFIER |
|---|---|---|
| Deposited data |  |  |
| TRFs in sensor and source space | Karl D. Lerud, Zenodo.org | https://doi.org/10.5281/zenodo.23773280 |
| Software and algorithms |  |  |
| Custom EEG and MEG TRF analysis code | Karl D. Lerud,<br>https://github.com/lerud/eegAndMegAnalysis/tree/releases | https://doi.org/10.5281/zenodo.23813431 |
| Harvard-Oxford volumetric Cortical Structural Atlas | FSL<sup>44,45</sup> | RRID:SCR_001476 |
| FreeSurfer<sup>56</sup> | MGH Harvard | RRID:SCR_001847 |
| MNE-Python<sup>42</sup> | MNE Developers | RRID:SCR_005972 |
| Eelbrain<sup>57</sup> | Christian Brodbeck | RRID:SCR_014661 |
| Presentation | Neurobehavioral systems,<br>https://www.neurobs.com | RRID:SCR_002521 |
| NiiVue<sup>58</sup> | NiiVue Developers | https://doi.org/10.5281/zenodo.18773294 |

### Experimental model and study participant details

#### Participants and EEG/MEG/MRI data acquisition

The 12 participants (8 females) were aged 18-29 years (mean 23 years) and were native English speakers; they were part of a larger study (in which the click train was not presented to all participants), the results of which will be presented separately. Pure tone thresholds were measured from 125 Hz to 8 kHz and all participants had thresholds less than 20 dB HL in at least one ear. This study included two sessions. In the first session, participants listened to competing speaker and click train stimuli with concurrent EEG and MEG recording. A second session included a more extensive behavioral experiment (not reported on here), and the structural MRI scan utilized for this analysis.

During the first session, a 32-channel BioSemi EEG cap was placed on all participants. An MEG system with 157 axial gradiometers (KIT-Eagle Technology, Kanazawa, Japan) was used to record the brain’s magnetic fields. Magnetic gradiometers are not especially sensitive to subcortical sources, but they are sensitive enough, for this dataset, to reveal robust wave V responses (Figure 1B, D) from auditory midbrain (Figure 4A). EEG was recorded at a rate of 16,384 Hz and the MEG was recorded at a rate of 2,000 Hz. The head shape, EEG electrode positions, and five head marker coils were digitized for each participant with a Polhemus 3SPACE FASTRAK three-dimensional digitizer and participant head positions were recorded in the MEG to facilitate forward models. Triggers were delivered to the EEG and MEG datasets during the experiment at the start and end of every trial digitally (through the Presentation® script (Neurobehavioral Systems, Inc., Berkeley, CA, www.neurobs.com) delivering the audio files) and in an analog fashion (with clicks at the start and end of the audio file in a non-audio channel, detected with an external Triggy hardware box (Cortech Solutions, Wilmington NC, USA).

At a separate session, T1-weighted structural MRI scans for each subject were acquired in the sagittal plane using a magnetization-prepared rapid gradient echo (MPRAGE) sequence on a Siemens Magnetom Prisma Fit (Siemens Healthcare, Erlangen, Germany) located at the Maryland Neuroimaging Center at the University of Maryland. The field strength was 3 Tesla (3T), slice thickness 0.8 mm (for 0.8 x 0.8 x 0.8 mm^3^ voxels), 2400 ms repetition time, 2 ms echo time, 1060 ms inversion time, and 8° flip angle.

#### Stimulus preparation and presentation

Speech passage stimuli were generated with Google text-to-speech synthesis^59^ utilizing four voices from their WaveNet model: British and American female and male. Parameters were passed to Google’s API, and then to Praat,^60^ to control the female fundamental frequencies to have a mean of 215 Hz, to control the male fundamental frequencies to have a mean of 110 Hz, and for all stimuli to minimize silence gaps. All stimuli were delivered at 70 dB SPL at the tympanic membrane calibrated with an acoustic level meter. Participants, lying in a supine position, heard audio stimuli diotically through foam eartips and 3.1-meter-long tubephones with Etymotic ER-3A transducers located outside the magnetically-shielded MEG room (Vacuumschmelze GmbH & Co. KG, Hanau, Germany) to prevent stimulus artifact.

Each participant listened to 32 1-minute speech trials in total, 24 of which had both a target and a distractor speaker, and 8 of which had only one speaker. The 8 trials with only a single speaker are not analyzed here to facilitate comparisons. For the two-speaker trials, target and distractor speakers were either different sexes or had different accents. Participants’ attention was directed to a particular speaker by telling them, and showing them with a picture, the sex (female or male) and accent (British or American English) of the speaker to attend to. A True/False comprehension question was asked at the end of each trial, to further encourage the participants to attend to the target. Target/distractor SNR varied from 4 to -4 dB but averaged to 0 dB; analysis of separate responses to the target and distractor speakers was not performed here in detail and will be presented in a separate analysis. However, a summary and plots of sensor-space target and distractor TRFs are provided for completeness in Figures S3 and S8.

The 10-minute click train consisted of unipolar clicks of 0.2 ms length, at an overall level of 70 dB SPL, and with inter-onset intervals (IOIs) sampled randomly from a distribution of the reciprocal of logarithmically-spaced frequencies between 5 Hz and 230 Hz. This distribution of IOIs spanned the range from slower IOIs common in evoked ABR-MLR experiments to faster IOIs comprising human speech fundamental frequencies (roughly 80 – 230 Hz). Sensor space EEG auditory evoked responses to clicks are shown in Figure S4. 200 50-ms 1 kHz tone pips were also randomly interspersed, which the subjects were given the task of counting in order to encourage attending to the stimulus, but the responses to the tones were not analyzed here.

## Method details

### Regressors and EEG/MEG processing

Regressors for the speech stimuli were created by taking the acoustic speech mixture and passing it to the auditory nerve model^35^ and saving the spike rates sampled at 100,000 Hz. The regressor for the click train was created analogously, by taking the clicks portion of the continuous stimulus (excluding the tone pips) and passing it to the auditory nerve model in the same way. The auditory nerve model implementation consists of 43 logarithmically spaced frequency channels with center frequencies from 125 Hz to 16,000 Hz. The model simulated high spontaneous rate auditory nerve fibers and included a fractal noise term. The 43 model outputs corresponding to these frequency channels were averaged to create a single, 1-dimensional regressor representing the entire spectrum. The model was run anew for each subject, including for different subjects in the same counterbalance group (thus identical stimuli), allowing subject-to-subject internal noise variation for more naturalistic simulation.

Additional processing was done to compensate for delays. The speed of sound inside the tubephones was compensated for in data processing by adding time to all event onsets: The tube length was 3.1 meters, and the speed of sound is 343 meters per second, therefore 3.1 / 343 = 0.00904 seconds (9.04 ms) was added to event onsets, shifting TRFs to the left (earlier) by 9.04 ms. 2.75 ms was also subtracted from event onsets to compensate for the auditory nerve delay built into the auditory nerve model utilized here.^34^ We found that the computer delivering the stimuli had a slightly faster clock rate relative to the recording computers, resulting in trials that were ∼6 ms shorter than the original minute-long speech audio stimuli that were delivered (and, analogously, click train trials that were ∼60 ms shorter than the 10-minute-long click audio stimuli). In order to be able to use the auditory nerve model regressors that were created from the audio files, we resampled the EEG and MEG datasets to correspond to the original audio files, thus correcting for the real-world, incorrect sampling rate that the stimuli were actually delivered at. This step is crucial to the ability to deconvolve high-frequency TRFs such as the ABR, as has been documented in previous recent work.^31,34^

MNE-Python^40,42^ was used for EEG and MEG processing. EEG was resampled from 16,384 Hz down to 5,000 Hz, and MEG was upsampled from 2,000 Hz to 5,000 Hz to match the EEG. The regressors were also downsampled to 5,000 Hz to match the EEG/MEG data sampling rate. EEG/MEG and all regressors were notch filtered with a minimum phase causal filter with a transition bandwidth of 1 Hz to remove all harmonics of 60 Hz. EEG/MEG and all regressors were also bandpass filtered between 20 and 300 Hz with a minimum phase causal filter to focus the responses on the part of the spectrum commonly understood to contain the ABR and MLR. As both data and regressors were filtered with the same filters, no further compensation for delays was necessary (beyond those discussed above to compensate for the speed of sound and the auditory nerve model intrinsic delay).

A TRF is generated from a continuous stimulus by deconvolving a stimulus-feature regressor from the continuous response data. TRFs were deconvolved concurrently for EEG and MEG in sensor space with MNE-Python using the mne.decoding.ReceptiveField class. A simple least-squares regression was used without regularization of any kind, corresponding to

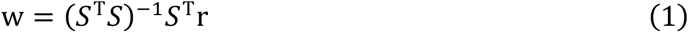

where *S* is the matrix of lagged stimulus regressor (auditory nerve model in this case) time series, *r* the matrix of EEG/MEG response data, and *w* the resultant matrix of TRFs per channel. Further, all speech and click TRFs were derived through complete *k*-fold cross validation with data splits every minute. Single-minute splits were natural for the speech experimental design, as each separate trial was one minute of speech. The total click train stimulus, which was 10 minutes long and consisted of 12,000 clicks, was split into 10 single-minute folds. As the TRF at each sensor then has an associated model goodness of fit, the corresponding product-moment correlations between the predicted and actual sensor data (thus the encoding model) are depicted in Figure S5.

To choose timings to visualize the topographies, as well as further analysis, latencies of the main ABR and MLR peaks were calculated algorithmically. This was done based on phase-locking between sensors within the expected ranges of peak occurrences and was calculated for each subject, each condition, and average TRFs over subjects, in the ranges of wave V, SN10, Pa, Nb, and P50. The resulting topographies are shown in Figure 1 for both EEG and MEG.

### Source-space localization and anatomical processing

Source space volumes and forward models were constructed utilizing each subject’s brain segmentation, skin and skull surfaces, and sensor locations, and inverse modeling resulted in a minimum norm solution using MNE-Python.^40–42^ Each T1 was segmented and parcellated using the FreeSurfer v7.4.1^56^ recon-all pipeline, and boundary element surfaces (scalp, outer skull, inner skull, brain) were constructed with the FreeSurfer watershed method using MNE-Python’s mne.bem.make_watershed_bem function. These surfaces, along with the individually digitized EEG electrode locations, the head position inside the MEG bore, and a homogenous brain volume source space, comprised the individual forward model for each participant. The volume source space used 8 mm^3^ vertex spacing.

Inverse modeling was done using a minimum norm solution and the dynamic statistical parametric mapping (dSPM) measure, with SNR of 9 and depth weighting 0.8, using MNE-Python; dSPM helps to control for the inherent bias the minimum norm solution has for superficial sources,^61^ and a higher assumed SNR corresponds to a lower level of regularization in the inverse model, allowing for a smaller point-spread function and more precise source localization.^62^ Point-spread functions were calculated for points (in an average brain) at the inferior colliculus, medial geniculate body, and right auditory cortex, and are plotted in Figure S1. Also, while the assumed SNR of 9 in the inverse model allows for more precise sources than the smaller values, we note that the present results are stable at other values, including the more standard value of 3 which is suggested as a default in most MNE documentation. The complete speech-minus-click TRF difference results for SNR=3 are in Table S1. The SN10, along with other clusters comparable with the SNR=9 results here, remains significant.

ROIs from the Destrieux atlas parcellation^46^ were used for individual ROI time series extraction (e.g. Figure 5), including Heschl’s gyrus and transverse temporal sulcus, brainstem, and bilateral thalamus. After completing source reconstruction for each participant, all source-space TRFs were morphed to a common head space—in this case, fsaverage which is FreeSurfer’s default average template. With all source-space TRFs then in a common space comprising 3,553 vertices sampling the entire brain, TRFs could be subjected to statistical comparisons and averaged over subjects for visualization and other analysis. TRFs were z-scored for each condition, and it was at this stage that spatiotemporal cluster permutation tests were performed.

### Ǫuantification and statistical analysis

To quantify any significant differences in sensor space between speech TRFs and click TRFs, we used one-sample spatiotemporal cluster permutation tests.^63,64^ With these tests, we analyzed EEG and MEG separately for the speech vs. click comparison. All sensor-space tests utilized a cluster-forming threshold based on a *t(11)* distribution (12 participants minus one degree of freedom) and *α* = 0.05, corresponding to *t* = 2.201, and a complete set of permutations, leading to an exact test.

Because of the significant difference found between the wave V latencies for speech and click TRFs, and the need to time-lock the subsequent MLR portions of the TRFs (10 – 60 ms) accurately to each other when testing for differences, for all further analysis we time-locked the averaged TRFs to their wave V latencies, for each participant and each stimulus type. The same spatiotemporal cluster permutation tests were then performed again in sensor space, for both modalities and conditions, after wave V locking.

To observe differences in source space, we tested for significant differences between each TRF and 0, and for significant differences between the speech and click TRFs. We again first synchronized the wave V latencies for each TRF, time-locking the source-space time series per participant and per condition to the wave V times before further source-space analysis. After transforming the sensor-space TRFs into the volume source spaces (maintaining the full vector-valued, three-dimensional neural current source solutions), we subjected the TRFs to a whole-brain TFCE cluster analysis^37^ to test for differences between each condition and zero (one-sample tests) and between conditions (paired-samples test). Hotelling’s T-squared statistic was used in order to simultaneously test all three Cartesian dimensions of the vector-valued solution^43^ as implemented in Eelbrain’s^57^ eelbrain.testnd.Vector class. While *p*-values for individual clusters from the TFCE test are already corrected, we employed a more conservative threshold by only considering corrected *p*-values below 0.02.

To further quantify the difference in speech and click TRFs at the SN10 time (illustrated in Figure 7B), we also extracted SN10 peak times from the auditory cortex ROIs directly. To do this, we took the minimum value during the SN10 significant difference cluster time (Table 2) for each participant, for both the speech and click TRFs, from the auditory cortex ROIs. This source time course was extracted from the bilateral Heschl’s gyrus and sulcus ROIs. As seen in Figure S10, a paired-samples *t*-test reveals a significant difference between speech and click TRFs at the SN10 latency using this method as well.

For visualization of orientation and location using NiiVue^58,65^ such as Figures 4, 6, and 7, source-space TRFs and TRF differences masked by significance were interpolated to 1mm^3^ resolution, exported as NIfTI images, morphed again to MNI152 average space to correspond to NiiVue’s default brain, and plotted. To visualize the moment-to-moment current dipole differences on the brain, we projected the three-dimensional vector-valued inverse solution to just two dimensions: superior-inferior and anterior-posterior (for these auditory processing stages of interest, visualizations of how the left-right dimension adds little information is shown in Figures S6 and S7). With a two-dimensional solution, we can then map the instantaneous current dipole orientation to a location on a circle by transforming from Cartesian coordinates to radians. Utilizing a circular colormap, we can plot the orientation of groups of vertices according to their angle on the unit circle in color on the brain (cf. Figure 4E, 6D, and 7C). Because current orientation information by itself would not preserve magnitude information, for visualization purposes only, we represented magnitude with the percentage of total vertices in the source space that we actually plotted. Thus, a cluster of larger magnitude also appears larger in its extent with this method, while a cluster of smaller magnitude appears smaller in extent.

## Supplemental Information

Document S1. Supplemental methods, Figures S1-S11, and Tables S1-S4.

## Supplementary materials

**Figure S1:**
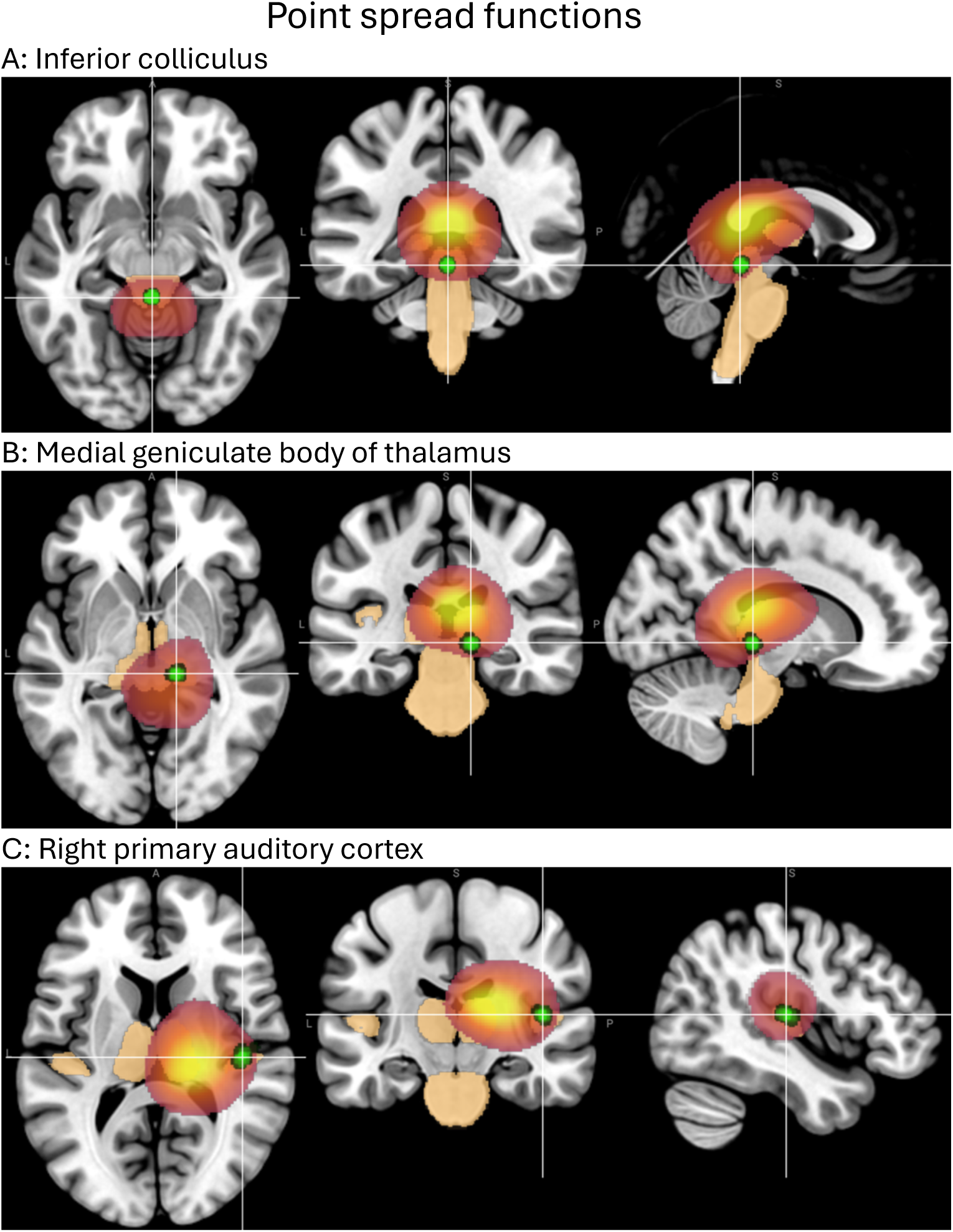
Point-spread functions (the spread of a source-localization solution of the inverse problem for a neural point source) for the (A) bilateral inferior colliculus (IC), (B) right medial geniculate body of the thalamus, and (C) right Heschl’s gyrus. As often occurs in minimum norm-based inverse solutions, the centroid of the point-spread function is not in general located at the point source itself.1 These functions clarify the main source-localization results, including the wave V TRF source location in e.g. Figure 4A, which, though appearing to be superior to the actual location of IC, is strongly consistent with how an IC source would manifest.

**Figure S2:**
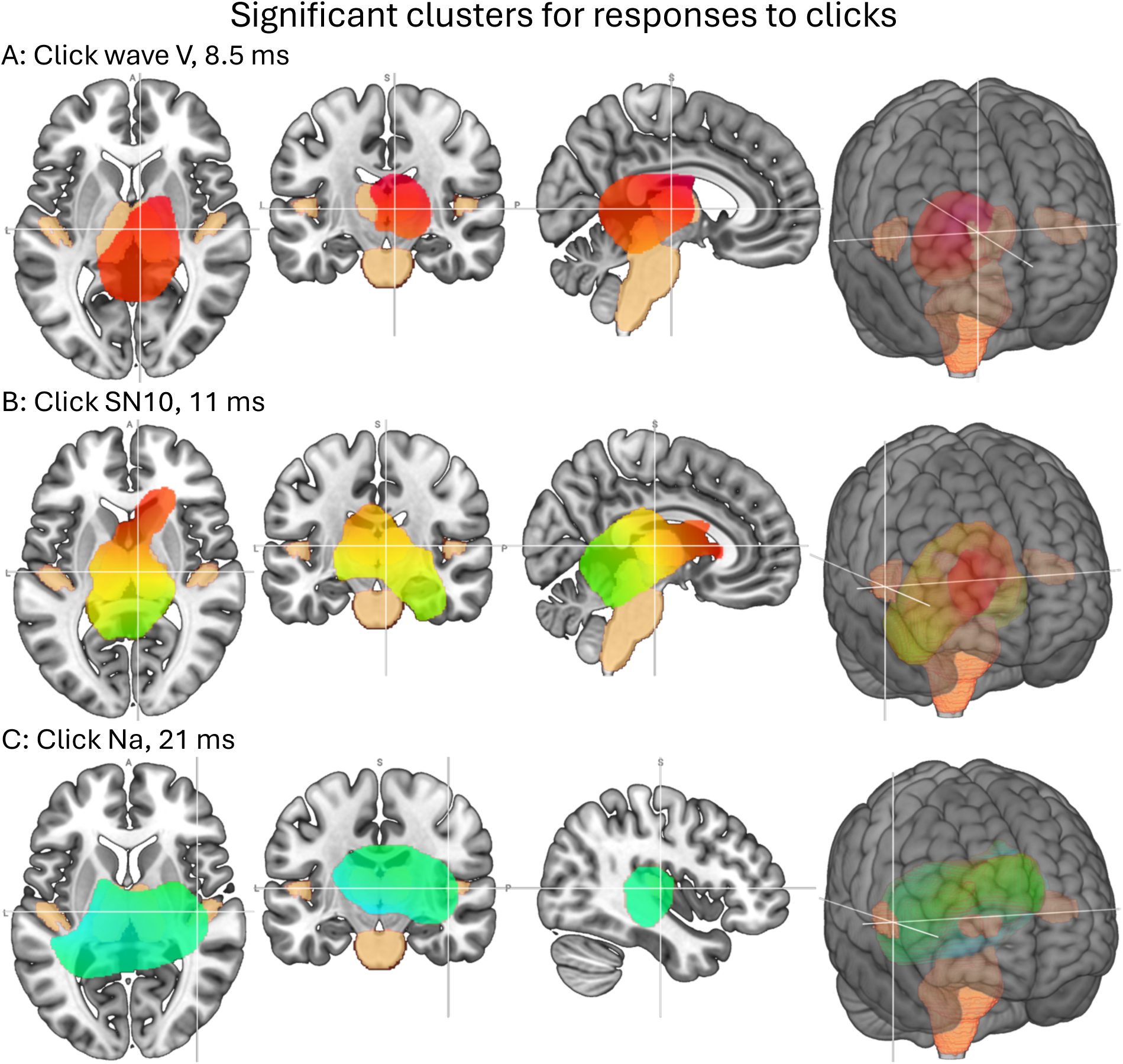
Early click TRF sources. Slice and render orientations as in Figure 4. Wave V (A) is nearly identical to wave V of the speech TRF (Figure 4A), while, in contrast, the distribution at the latency of the speech SN10 (B) shows little evidence of cortical sources. The Na (C) also shows weaker evidence of cortical activity than the comparable speech TRF response in Figure 4C.

**Figure S3:**
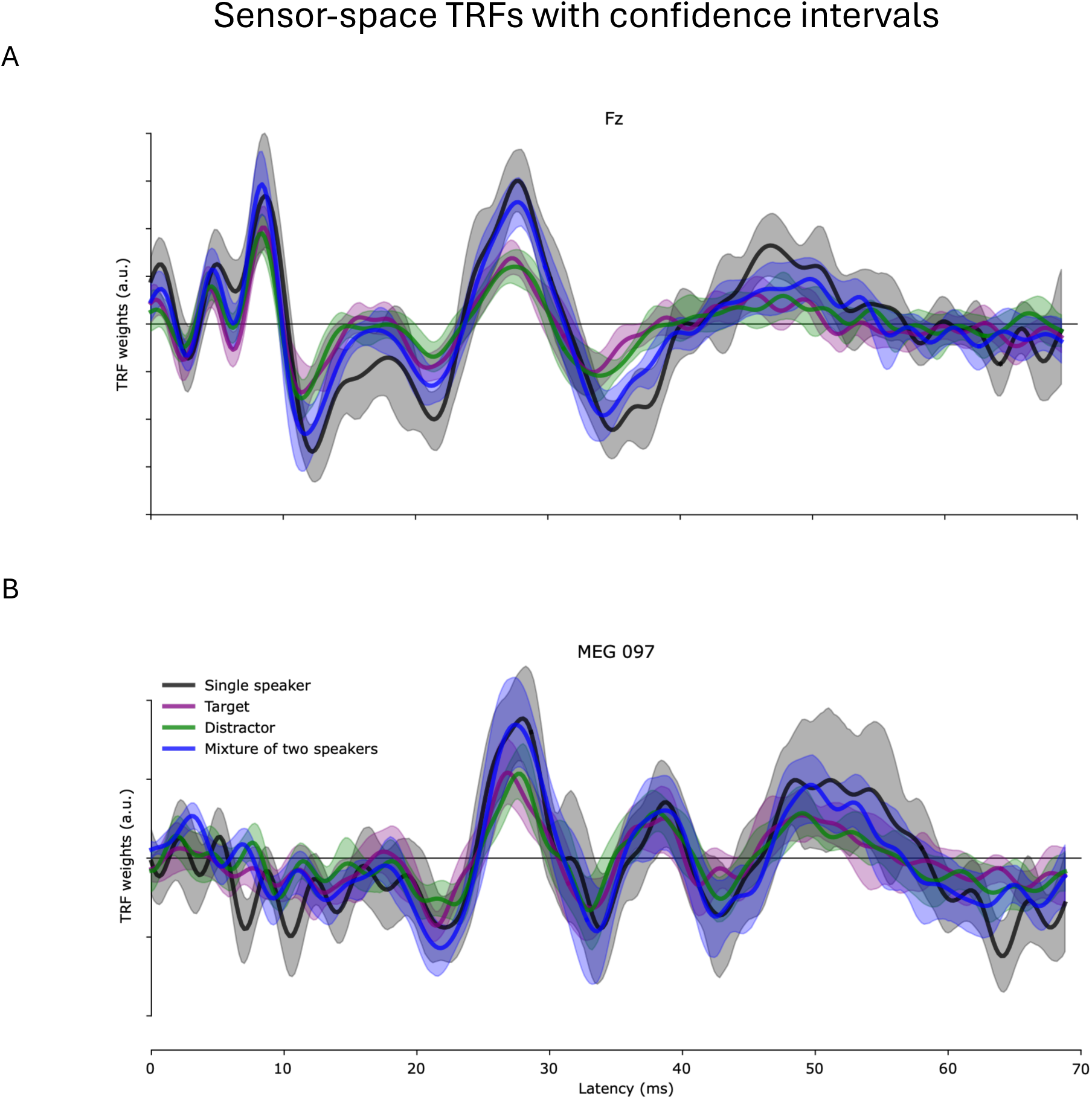
Representative single-sensor TRFs with 95% confidence intervals: (A) EEG, (B) MEG. Four TRFs shown: single speaker (speech in quiet) stimuli, target speaker analyzed separately from concurrent distractor, distractor speaker analyzed separately from concurrent target, and the full speech mixture (as used in this study’s main analysis).

**Figure S4:**
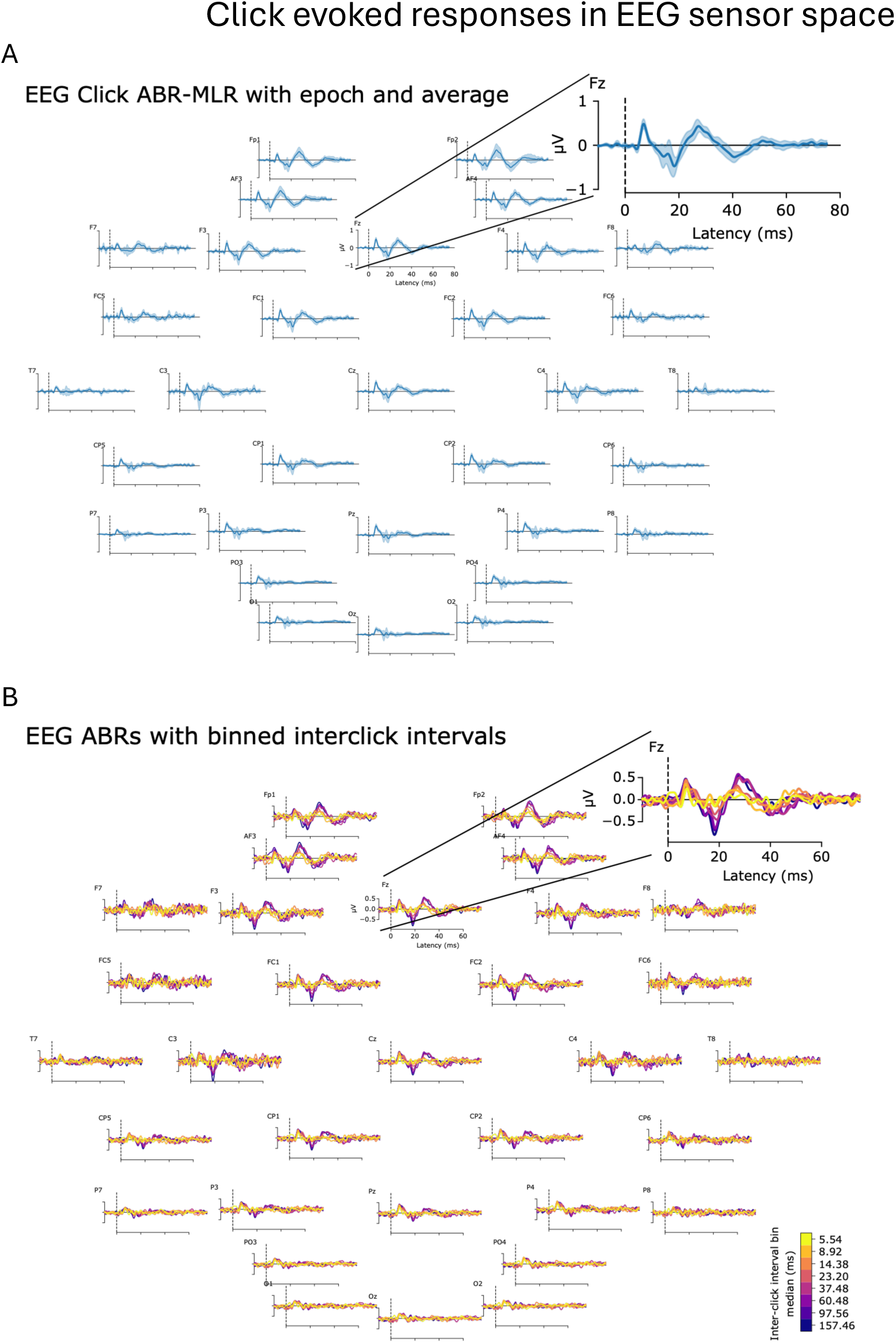
EEG sensor-space click evoked responses from epoching and averaging (in contrast to the TRFs shown elsewhere), (A) with 95% confidence intervals, and (B) separated into interclick interval bins (time since previous click).

**Figure S5:**
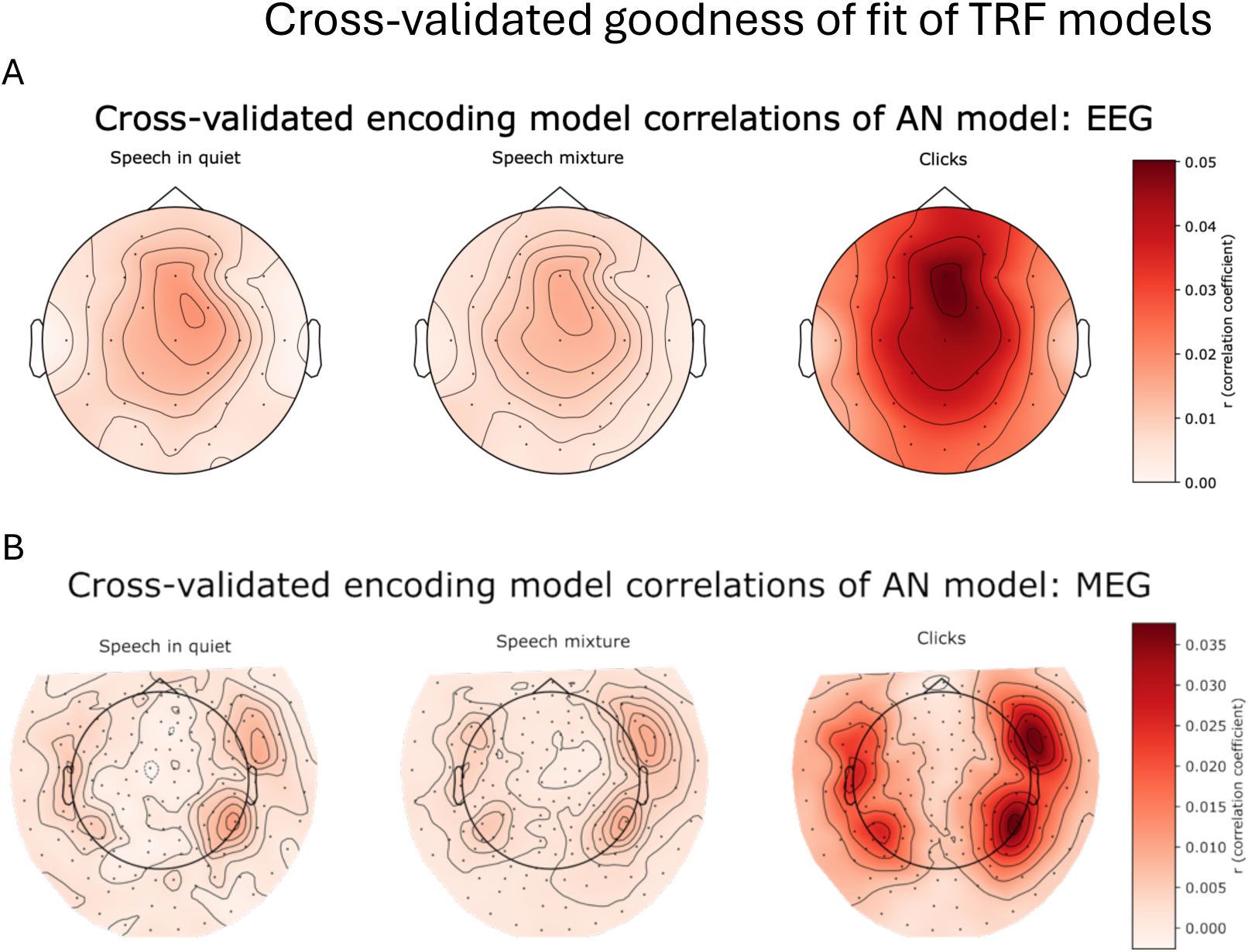
Distribution of cross-validated encoding model prediction accuracies of unseen data in sensor space for (A) EEG and (B) MEG.

**Figure S6:**
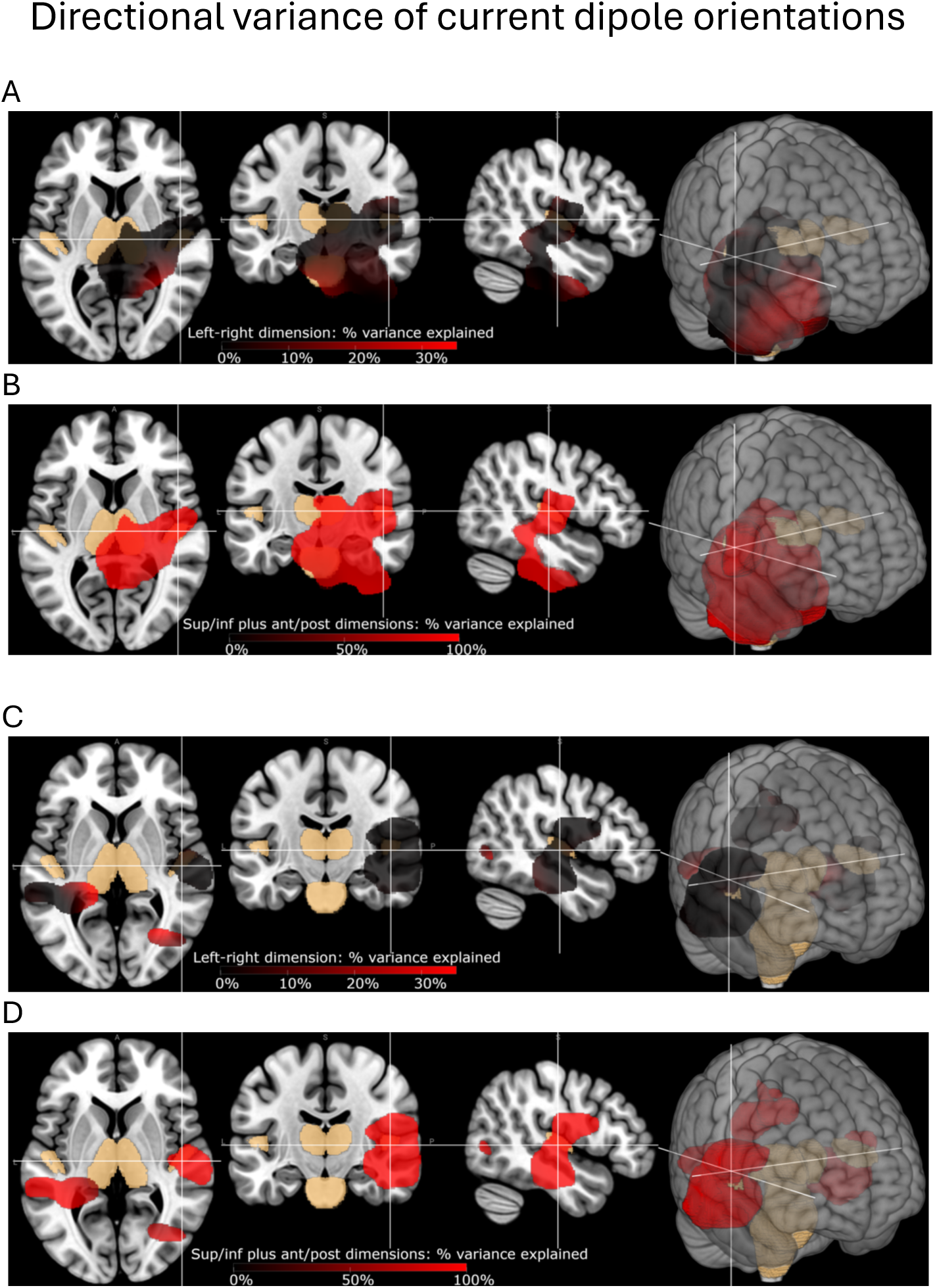
Supporting analysis for the (graphical-only) omission of the left-right-axis vector projections of current dipole orientations in Figures 4, C, 7, and S2. Percentage of directional variances explained by left-right dimension vs. combination of superior-inferior and anterior-posterior dimensions. Note that for the left-right dimension depictions (A and C), the colormap only goes as high as 33% to enhance detail. As can be seen, the left-right dimension contains little explained directional variance for the current dipole source orientations for the SN10 (A, B) and P50 (C, D) as examples. For this reason, visually omitting this dimension in Figures 4, 6, 7, and S2 suppresses only minimal source information in these particular auditory responses.

**Figure S7:**
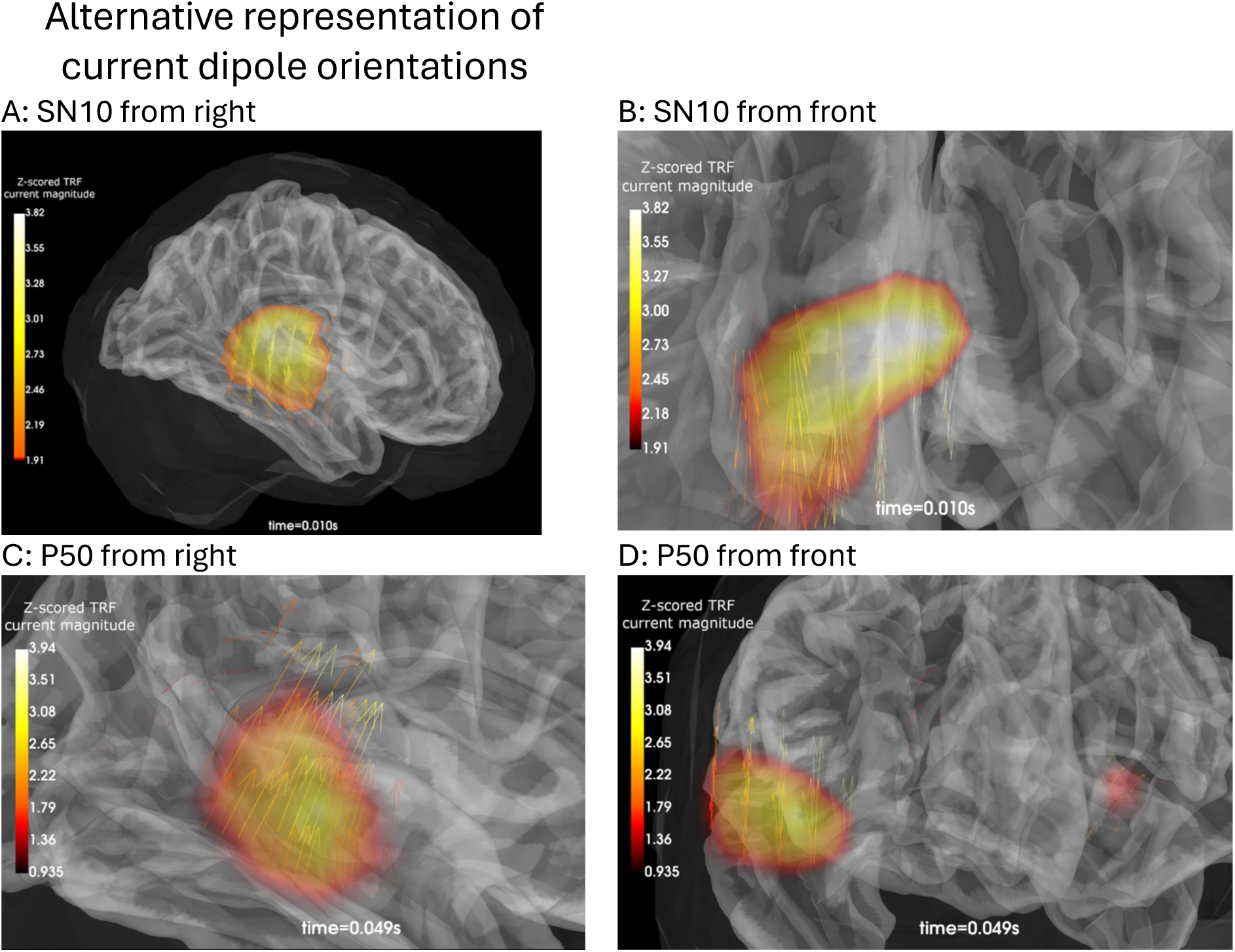
An alternative to using a color wheel to indicate current dipole direction is a more traditional vector field representation, here the quiver/arrow representations from MNE-Python. (A) and (B) show the SN10 from the right and front vantage points, respectively. (C) and (D) show the P50, also from the right and front vantage points. It can be seen from (B) and (D) that the left-right vector component of the solution is minimal compared to the other two dimensions, as is also depicted in Figure SC.

**Figure S8:**
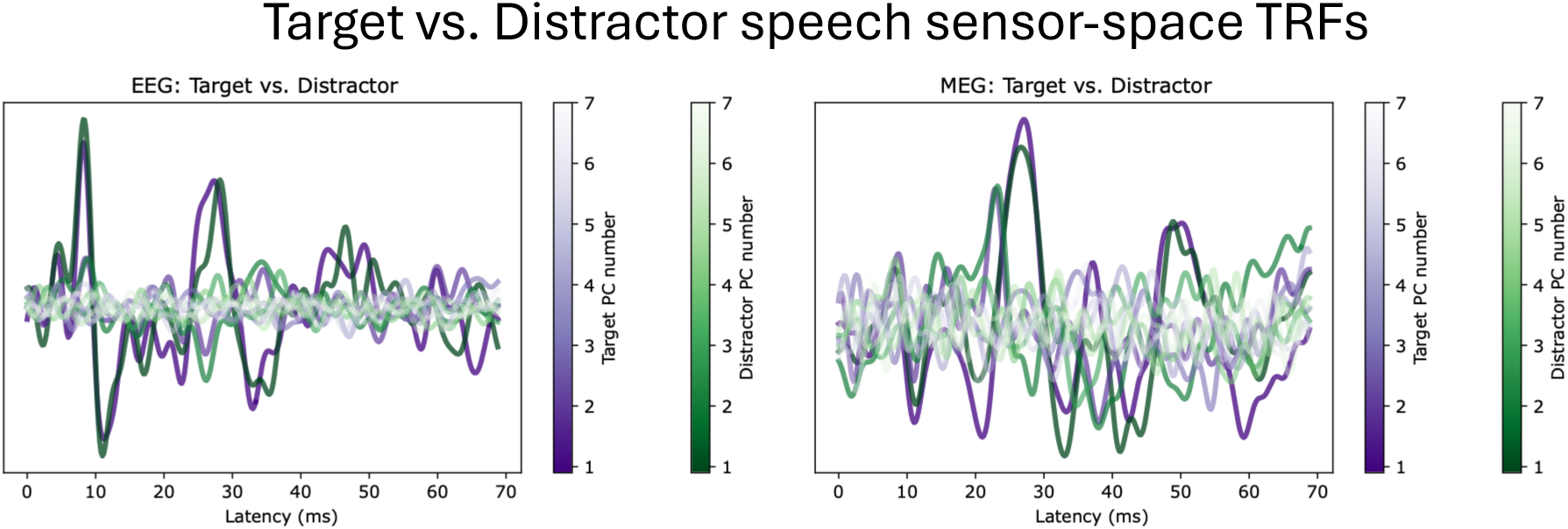
Target and distractor TRFs in sensor-space principal component space, in the style of Figure 2. Sensor-space and source-space spatiotemporal cluster tests did not show a significant diaerence between these two TRFs in this study. Increasingly lighter shades of purple and green represent higher-order components of PCA, from number 1 through 7, for the target and distractor TRFs, respectively.

**Figure S9:**
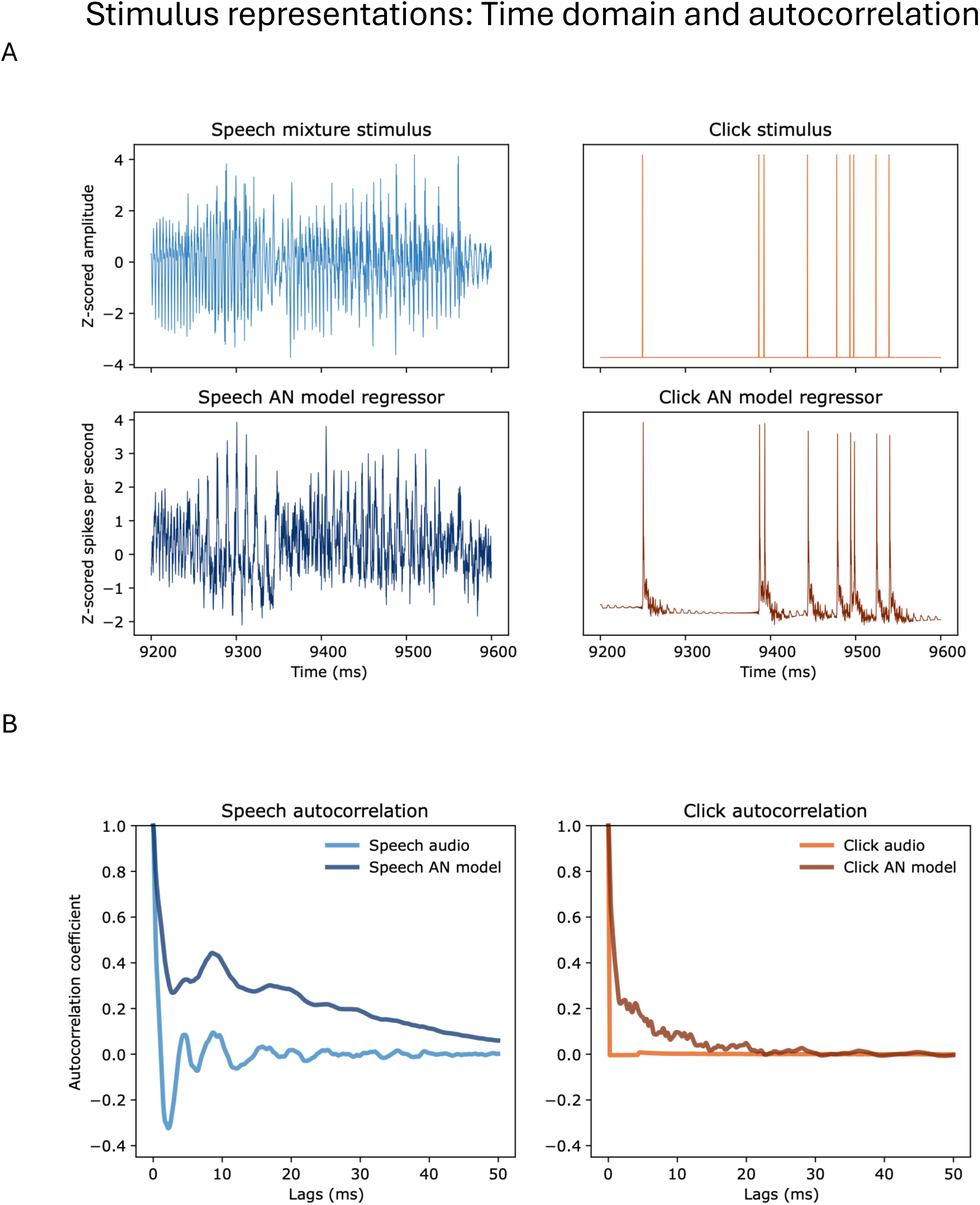
Example stimulus acoustic and regressor representations in the time domain and as autocorrelation functions. (A) Time domain representations show only 400 ms of time to enhance detail; (B) autocorrelations are calculated over the entire length of the corresponding speech and click stimuli. Speech shows characteristic peaks in its autocorrelation corresponding to average fundamental frequencies of female (200 Hz, 5 ms) and male (100 Hz, 10 ms) speech. The click train acoustic representation does not show any autocorrelation, consistent with its generation from randomly sampled interclick intervals.

**Figure S10:**
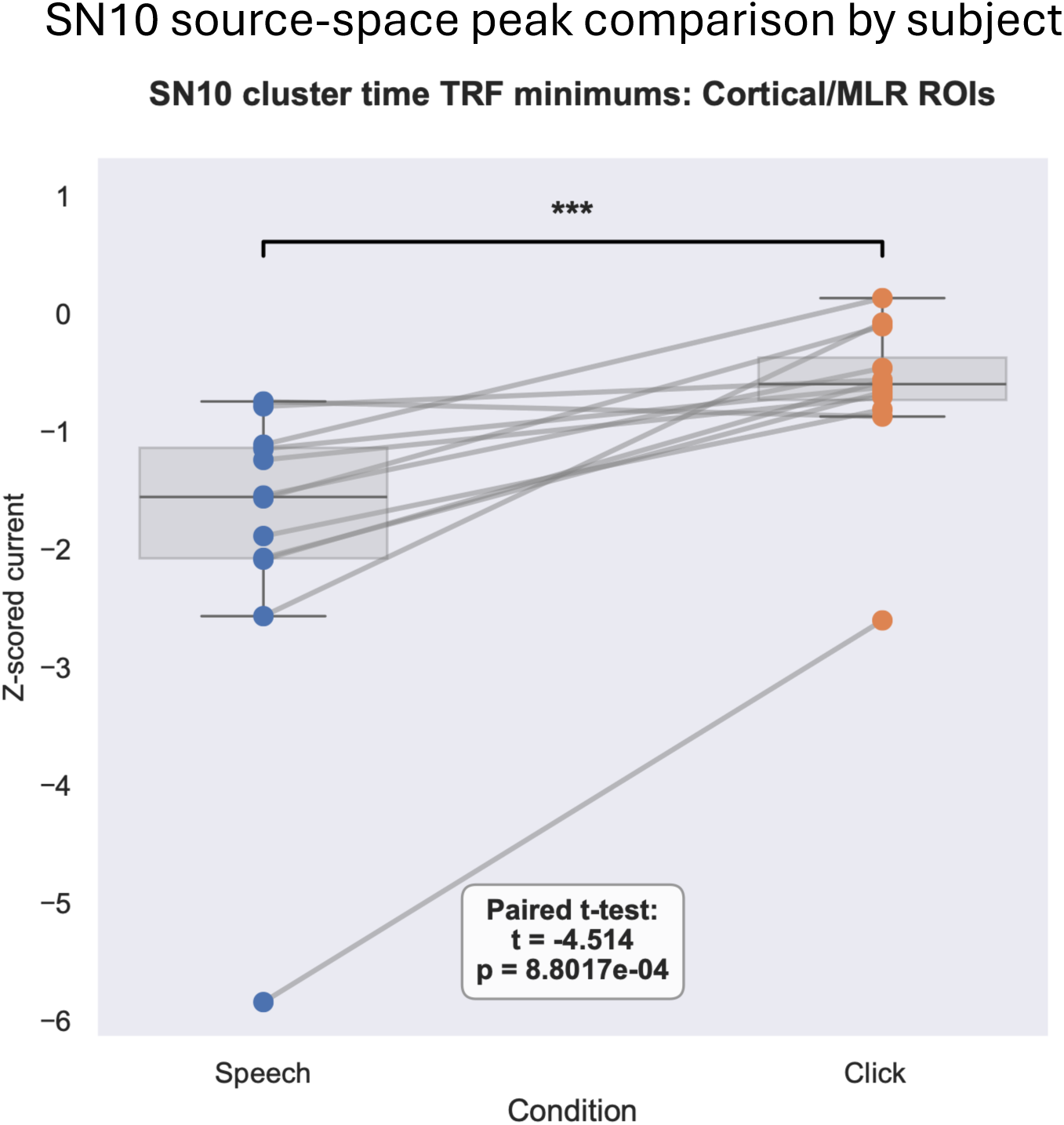
Paired-samples t-test on SN10 source-space TRF current weight from auditory cortex ROIs. This corresponds to the cluster result at the SN10 time showing a significant difference between the speech and click TRFs, re-illustrating that the auditory cortical source space TRF has a more negative peak at the SN10 latency for speech than for clicks.

**Figure S11:**
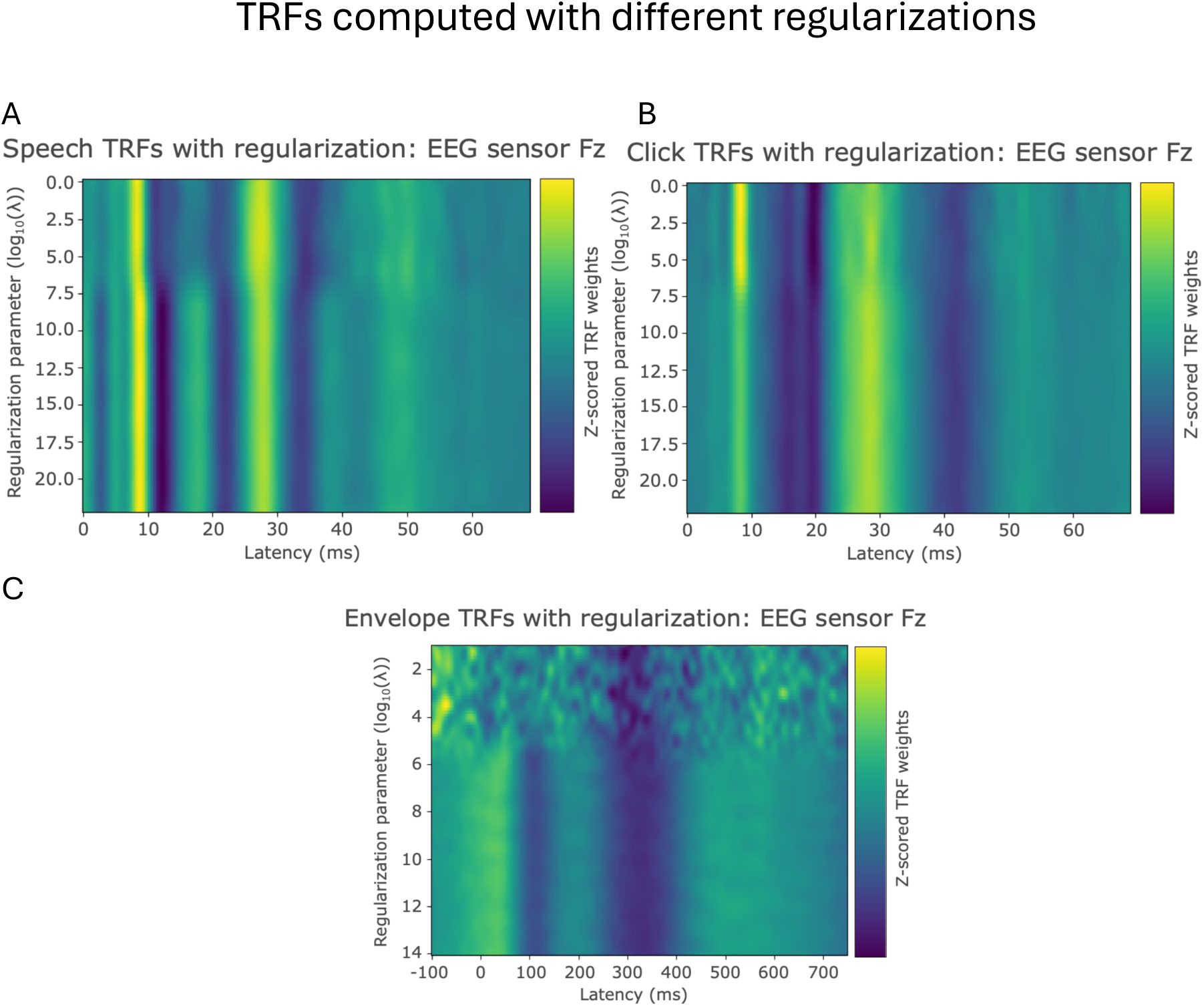
TRF weights as a function of regularization parameter lambda. Peak times for the speech TRF (A) shift slightly with increasing regularization, while click TRF peaks (B) mostly do not. This may indicate that the respective autocorrelation functions leak more strongly (respectively, less strongly) into the TRF’s morphology when computed with regularized ridge regression. Note also that both show clean TRF morphologies at all values of the regularization parameter, including 0 (the topmost row). In contrast, (C), an example lower-frequency (stimulus representations and responses low-passed at 40 Hz) auditory envelope TRF, demonstrates that at low frequencies regularization is clearly necessary, as an insufficient amount produces TRFs that only fit the noise.

**Table S1:** Source-space cluster statistics for SNR=3. All main results comparable to SNR=9 in the main analysis, including the SN10 time (9.6 to 12.6 ms here) are present.

| *# of source vertices* | *Start time (ms)* | *Stop time (ms)* | p-*value* |
|---|---|---|---|
| 2472 | 38.4 | 59.6 | 0.0004884 |
| 2405 | 23.8 | 36.6 | 0.0004884 |
| 342 | 9.6 | 12.6 | 0.0063492 |
| 175 | 16.4 | 19.8 | 0.008547 |
| 373 | 56.6 | 62.2 | 0.016117 |
| 16 | 42.4 | 44 | 0.027595 |
| 119 | 52.8 | 54.8 | 0.027839 |
| 32 | 13.8 | 15 | 0.033455 |
| 55 | 59.8 | 61.2 | 0.035653 |
| 10 | 41.2 | 41.8 | 0.042247 |
| 40 | 50.2 | 51.4 | 0.042491 |
| 14 | 4.6 | 5.2 | 0.042979 |
| 12 | 52.4 | 54 | 0.043223 |
| 12 | 18.8 | 20.4 | 0.044933 |
| 7 | 68 | 68.6 | 0.046642 |
| 1 | 6.2 | 6.4 | 0.047375 |
| 1 | 18.8 | 19.4 | 0.048107 |
| 6 | 68.6 | 69 | 0.048107 |
| 5 | 58.8 | 59.6 | 0.048107 |
| 4 | 42.2 | 42.8 | 0.048107 |
| 1 | 19 | 19.2 | 0.049817 |

**Table S2:** Interhemispheric test for significance for speech minus click difference. Hemispheres for each participant were reversed, and the test was non-reversed minus reversed speech minus click TRF difference source images. The difference is significant, with polarity of clusters in the right hemisphere being the same polarity as the original difference image. As the paired left-hemisphere clusters are at opposite polarity, this indicates that there was significantly more right-hemisphere activity in the speech minus click difference. There is no cluster at the SN10 time, however, so the apparent right hemisphere dominance of vertices of the SN10 cluster from the original speech minus click difference test (Figure 7B) are not significant.

| *# of source vertices* | *Start time (ms)* | *Stop time (ms)* | p-*value* |
|---|---|---|---|
| 36 | 39 | 40.2 | 0.013431 |
| 36 | 39 | 40.2 | 0.013431 |
| 18 | 31.2 | 32.8 | 0.01978 |
| 2 | 38.4 | 39 | 0.024908 |
| 16 | 32.4 | 33.2 | 0.026618 |
| 16 | 32.4 | 33.2 | 0.026618 |
| 2 | 38.4 | 39 | 0.026618 |
| 10 | 33.2 | 34.6 | 0.037851 |
| 5 | 50.2 | 50.8 | 0.04127 |
| 5 | 50.2 | 50.8 | 0.04127 |

**Table S3:** Source-space cluster statistics for speech TRF difference from zero.

| *# of source vertices* | *Start time (ms)* | *Stop time (ms)* | p-*value* |
|---|---|---|---|
| *2215* | 16 | 42 | 0.0002 |
| *1649* | 3.2 | 16.4 | 0.002 |
| *117* | 40 | 44 | 0.0054 |
| *140* | 1 | 3.6 | 0.0086 |
| *43* | 42.6 | 43.6 | 0.0294 |
| *1* | 48.4 | 49 | 0.0365 |
| *21* | 48.6 | 49.6 | 0.0374 |
| *2* | 50.2 | 50.6 | 0.0395 |
| *11* | 51.6 | 52.2 | 0.0413 |
| *11* | 49.8 | 51 | 0.042 |
| *1* | 18.2 | 18.8 | 0.0455 |
| *1* | 18.2 | 18.4 | 0.0492 |
| *1* | 40 | 40.2 | 0.0497 |
| *1* | 20.4 | 20.6 | 0.05 |

**Table S4:** Source-space cluster statistics for click TRF difference from zero.

| *# of source vertices* | *Start time (ms)* | *Stop time (ms)* | p-*value* |
|---|---|---|---|
| 3512 | 0 | 64.2 | 0 |
| 1 | 1 | 1.2 | 0.0491 |

## References

1. Akram S, Simon JZ, Babadi B. Dynamic Estimation of the Auditory Temporal Response Function From MEG in Competing-Speaker Environments. IEEE Trans Biomed Eng. 2017;64(8):1896–1905. doi:10.1109/TBME.2016.2628884

2. Brodbeck C, Hong LE, Simon JZ. Rapid Transformation from Auditory to Linguistic Representations of Continuous Speech. Curr Biol. 2018;28(24):3976–3983.e5. doi:10.1016/j.cub.2018.10.042

3. Brodbeck C, Jiao A, Hong LE, Simon JZ. Neural speech restoration at the cocktail party: Auditory cortex recovers masked speech of both attended and ignored speakers. PLOS Biol. 2020;18(10):e3000883. doi:10.1371/journal.pbio.3000883

4. Ding N, Simon JZ. Emergence of neural encoding of auditory objects while listening to competing speakers. Proc Natl Acad Sci. 2012;109(29):11854–11859. doi:10.1073/pnas.1205381109

5. O’Sullivan J, Herrero J, Smith E, et al. Hierarchical Encoding of Attended Auditory Objects in Multi-talker Speech Perception. Neuron. 2019;104(6):1195–1209.e3. doi:10.1016/j.neuron.2019.09.007

6. O’Sullivan JA, Power AJ, Mesgarani N, et al. Attentional Selection in a Cocktail Party Environment Can Be Decoded from Single-Trial EEG. Cereb Cortex. 2015;25(7):1697–1706. doi:10.1093/cercor/bht355

7. Power AJ, Foxe JJ, Forde EJ, Reilly RB, Lalor EC. At what time is the cocktail party? A late locus of selective attention to natural speech. Eur J Neurosci. 2012;35(9):1497–1503. doi:10.1111/j.1460-9568.2012.08060.x

8. Coffey EBJ, Herholz SC, Chepesiuk AMP, Baillet S, Zatorre RJ. Cortical contributions to the auditory frequency-following response revealed by MEG. Nat Commun. 2016;7:1–11. doi:10.1038/ncomms11070

9. Kulasingham JP, Brodbeck C, Presacco A, Kuchinsky SE, Anderson S, Simon JZ. High gamma cortical processing of continuous speech in younger and older listeners. NeuroImage. 2020;222(June):117291. doi:10.1016/j.neuroimage.2020.117291

10. Schüller A, Schilling A, Krauss P, Rampp S, Reichenbach T. Attentional Modulation of the Cortical Contribution to the Frequency-Following Response Evoked by Continuous Speech. J Neurosci. 2023;43(44):7429–7440. doi:10.1523/JNEUROSCI.1247-23.2023

11. Schüller A, Schilling A, Krauss P, Reichenbach T. The Early Subcortical Response at the Fundamental Frequency of Speech Is Temporally Separated from Later Cortical Contributions. J Cogn Neurosci. 2024;36(3):475–491. doi:10.1162/jocn_a_02103

12. Land R, Burghard A, Kral A. The contribution of inferior colliculus activity to the auditory brainstem response (ABR) in mice. Hear Res. 2016;341:109–118. doi:10.1016/j.heares.2016.08.008

13. Møller A, Jannetta P. Evoked potentials from the inferior colliculus in man. Electroencephalogr Clin Neurophysiol. 1982;53:612–620.

14. McGee T, Kraus N. Auditory Development Reflected by Middle Latency Response. Ear Hear. 1996;17(5):419–429. doi:10.1097/00003446-199610000-00008

15. Musiek F, Nagle S. The Middle Latency Response: A Review of Findings in Various Central Nervous System Lesions. J Am Acad Audiol. 2018;29(9):855–867. doi:10.3766/jaaa.16141

16. Burkard RF, Sims D. The Human Auditory Brainstem Response to High Click Rates. Am J Audiol. 2001;10(2):53–61. doi:10.1044/1059-0889(2001/008)

17. Delgado RE, Ozdamar O. Deconvolution of evoked responses obtained at high stimulus rates. J Acoust Soc Am. 2004;115(3):1242–1251. doi:10.1121/1.1639327

18. Don M, Allen AR, Starr A. Effect of Click Rate on the Latency of Auditory Brain Stem Responses in Humans. Ann Otol Rhinol Laryngol. 1977;86(2):186–195. doi:10.1177/000348947708600209

19. Serpanos YC, O’Malley H, Gravel JS. The Relationship between Loudness Intensity Functions and the Click-ABR Wave V Latency. Ear Hear. 1997;18(5):409.

20. Tucker DA, Dietrich S, Harris S, Pelletier S. Effects of Stimulus Rate and Gender on the Auditory Middle Latency Response. J Am Acad Audiol. 2002;13(3):146–153.

21. Tawfik S, Musiek FE. SN10 auditory evoked potential revisited. Am J Otol. 1991;12(3):179–183.

22. Dykstra AR, Burchard D, Starzynski C, Riedel H, Rupp A, Gutschalk A. Lateralization and Binaural Interaction of Middle-Latency and Late-Brainstem Components of the Auditory Evoked Response. J Assoc Res Otolaryngol. 2016;17(4):357–370. doi:10.1007/s10162-016-0572-x

23. Hashimoto I, Mashiko T, Yoshikawa K, Mizuta T, Imada T, Hayashi M. Neuromagnetic measurements of the human primary auditory response. Electroencephalogr Clin Neurophysiol. 1995;96(4):348–356. doi:10.1016/0168-5597(95)00004-C

24. Kuriki S, Nogai T, Hirata Y. Cortical sources of middle latency responses of auditory evoked magnetic field. Hear Res. 1995;92(1-2):47–51. doi:10.1016/0378-5955(95)00195-6

25. Mäkelä JP, Hämäläinen M, Hari R, McEvoy L. Whole-head mapping of middle-latency auditory evoked magnetic fields. Electroencephalogr Clin Neurophysiol Potentials Sect. 1994;92(5):414–421. doi:10.1016/0168-5597(94)90018-3

26. Brodbeck C, Simon JZ. Continuous speech processing. Curr Opin Physiol. 2020;18:25–31. doi:10.1016/j.cophys.2020.07.014

27. Ding N, Simon JZ. Neural coding of continuous speech in auditory cortex during monaural and dichotic listening. J Neurophysiol. 2012;107(1):78–89. doi:10.1152/jn.00297.2011

28. Lalor EC, Pearlmutter BA, Reilly RB, McDarby G, Foxe JJ. The VESPA: A method for the rapid estimation of a visual evoked potential. NeuroImage. 2006;32(4):1549–1561. doi:10.1016/j.neuroimage.2006.05.054

29. Lalor EC, Foxe JJ. Neural responses to uninterrupted natural speech can be extracted with precise temporal resolution. Eur J Neurosci. 2010;31(1):189–193. doi:10.1111/j.1460-9568.2009.07055.x

30. Bachmann FL, MacDonald EN, Hjortkjær J. Neural Measures of Pitch Processing in EEG Responses to Running Speech. Front Neurosci. 2021;15:738408. doi:10.3389/fnins.2021.738408

31. Bachmann FL, Kulasingham JP, Eskelund K, Enqvist M, Alickovic E, Innes-Brown H. Extending Subcortical EEG Responses to Continuous Speech to the Sound-Field. Trends Hear. 2024;28:23312165241246596. doi:10.1177/23312165241246596

32. Maddox RK, Lee AKC. Auditory Brainstem Responses to Continuous Natural Speech in Human Listeners. eNeuro. 2018;5(1). doi:10.1523/ENEURO.0441-17.2018

33. Polonenko MJ, Maddox RK. Exposing distinct subcortical components of the auditory brainstem response evoked by continuous naturalistic speech. eLife. 2021;10:e62329. doi:10.7554/eLife.62329

34. Shan T, Cappelloni MS, Maddox RK. Subcortical responses to music and speech are alike while cortical responses diverge. Sci Rep. 2024;14(1):1. doi:10.1038/s41598-023-50438-0

35. Zilany MSA, Bruce IC, Carney LH. Updated parameters and expanded simulation options for a model of the auditory periphery. J Acoust Soc Am. 2014;135(1):283–286. doi:10.1121/1.4837815

36. Kulasingham JP, Bachmann FL, Eskelund K, Enqvist M, Innes-Brown H, Alickovic E. Predictors for estimating subcortical EEG responses to continuous speech. PLOS ONE. 2024;19(2):e0297826. doi:10.1371/journal.pone.0297826

37. Smith SM, Nichols TE. Threshold-free cluster enhancement: Addressing problems of smoothing, threshold dependence and localisation in cluster inference. NeuroImage. 2009;44(1):83–98. doi:10.1016/j.neuroimage.2008.03.061

38. Zilany MSA, Bruce IC, Nelson PC, Carney LH. A phenomenological model of the synapse between the inner hair cell and auditory nerve: long-term adaptation with power-law dynamics. J Acoust Soc Am. 2009;126(5):2390–2412. doi:10.1121/1.3238250

39. Chait M, Poeppel D, Simon JZ. Neural Response Correlates of Detection of Monaurally and Binaurally Created Pitches in Humans. Cereb Cortex. 2006;16(6):835–848. doi:10.1093/cercor/bhj027

40. Gramfort A, Luessi M, Larson E, et al. MNE software for processing MEG and EEG data. NeuroImage. 2014;86:446–460. doi:10.1016/j.neuroimage.2013.10.027

41. Hämäläinen MS, Ilmoniemi RJ. Interpreting magnetic fields of the brain: minimum norm estimates. Med Biol Eng Comput. 1994;32(1):35–42. doi:10.1007/BF02512476

42. Larson E, Gramfort A, Engemann DA, et al. MNE-python. Published online November 2025. doi:10.5281/zenodo.17675410

43. Das P, Brodbeck C, Simon JZ, Babadi B. Neuro-current response functions: A unified approach to MEG source analysis under the continuous stimuli paradigm. NeuroImage. 2020;211:116528. doi:10.1016/j.neuroimage.2020.116528

44. Jenkinson M, Beckmann CF, Behrens TEJ, Woolrich MW, Smith SM. FSL. NeuroImage. 2012;62(2):782–790. doi:10.1016/j.neuroimage.2011.09.015

45. Smith SM, Jenkinson M, Woolrich MW, et al. Advances in functional and structural MR image analysis and implementation as FSL. NeuroImage. 2004;23:S208–S219. doi:10.1016/j.neuroimage.2004.07.051

46. Destrieux C, Fischl B, Dale A, Halgren E. Automatic parcellation of human cortical gyri and sulci using standard anatomical nomenclature. NeuroImage. 2010;53(1):1–15. doi:10.1016/j.neuroimage.2010.06.010

47. Parkkonen L, Fujiki N, Mäkelä JP. Sources of auditory brainstem responses revisited: Contribution by magnetoencephalography. Hum Brain Mapp. 2009;30(6):1772–1782. doi:10.1002/hbm.20788

48. Hamilton LS, Edwards E, Chang EF. A Spatial Map of Onset and Sustained Responses to Speech in the Human Superior Temporal Gyrus. Curr Biol. 2018;28(12):1860–1871.e4. doi:10.1016/j.cub.2018.04.033

49. Garcia MM, Kline AM, Onodera K, et al. Noncanonical short-latency auditory pathway directly activates deep cortical layers. Nat Commun. 2025;16(1):5911. doi:10.1038/s41467-025-61020-9

50. Liegeois-Chauvel C, Musolino A, Chauvel P. Localization of the primary auditory area in man. Brain. 1991;114(1):139–153. doi:10.1093/oxfordjournals.brain.a101854

51. Nourski KV, Steinschneider M, McMurray B, et al. Functional organization of human auditory cortex: Investigation of response latencies through direct recordings. NeuroImage. 2014;101:598–609. doi:10.1016/j.neuroimage.2014.07.004

52. Xie Z. Subcortical responses to continuous speech under bimodal divided attention. J Neurophysiol. Published online March 18, 2025. doi:10.1152/jn.00039.2025

53. Theunissen FE, David SV, Singh NC, Hsu A, Vinje WE, Gallant JL. Estimating spatio-temporal receptive fields of auditory and visual neurons from their responses to natural stimuli. Netw Comput Neural Syst. 2001;12(3):289. doi:10.1088/0954-898X/12/3/304

54. Etard O, Kegler M, Braiman C, Forte AE, Reichenbach T. Decoding of selective attention to continuous speech from the human auditory brainstem response. NeuroImage. 2019;200:1–11. doi:10.1016/j.neuroimage.2019.06.029

55. Forte AE, Etard O, Reichenbach T. The human auditory brainstem response to running speech reveals a subcortical mechanism for selective attention. Shinn-Cunningham BG, ed. eLife. 2017;6:e27203. doi:10.7554/eLife.27203

56. Fischl B. FreeSurfer. NeuroImage. 2012;62(2):774–781. doi:10.1016/j.neuroimage.2012.01.021

57. Brodbeck C, Das P, Gillis M, et al. Eelbrain, a Python toolkit for time-continuous analysis with temporal response functions. Martin AE, Shinn-Cunningham BG, Slaats S, eds. eLife. 2023;12:e85012. doi:10.7554/eLife.85012

58. Eckstein K, Androulakis A, Dao TT, et al. Seamless neuroimaging visualization: The NiiVue wrapper ecosystem. Aperture Neuro. 2026;6. doi:10.52294/001c.167815

59. Google. Google Cloud Text-to-Speech. Published online 2018. https://cloud.google.com/text-to-speech

60. Boersma P, Weenink W. Praat: doing phonetics by computer. Published online 2026. https://praat.org

61. Dale AM, Liu AK, Fischl BR, et al. Dynamic Statistical Parametric Mapping: Combining fMRI and MEG for High-Resolution Imaging of Cortical Activity. Neuron. 2000;26:55–67.

62. Hauk O, Stenroos M, Treder MS. EEG/MEG source estimation and spatial filtering: the linear toolkit. In: Magnetoencephalography: From Signals to Dynamic Cortical Networks. Springer International Publishing; 2019:167–203. doi:10.1007/978-3-030-00087-5

63. Maris E, Oostenveld R. Nonparametric statistical testing of EEG- and MEG-data. J Neurosci Methods. 2007;164(1):177–190. doi:10.1016/j.jneumeth.2007.03.024

64. Sassenhagen J, Draschkow D. Cluster-based permutation tests of MEG/EEG data do not establish significance of effect latency or location. Psychophysiology. 2019;56(6):e13335. doi:10.1111/psyp.13335

65. Hanayik T, Rorden C, Drake C, et al. niivue/niivue: @niivue/niivue-v0.68.1. Published online February 25, 2026. doi:10.5281/zenodo.18773294

## Supplementary References

1. Tait L, Özkan A, Szul MJ, Zhang J. A systematic evaluation of source reconstruction of resting MEG of the human brain with a new high-resolution atlas: Performance, precision, and parcellation. Hum Brain Mapp. 2021;42(14):4685–4707. doi:10.1002/hbm.25578

